# Spatially-resolved single cell transcriptomics reveal a critical role for γδ T cells in the control of skin inflammation and subcutaneous adipose wasting during chronic *Trypanosoma brucei* infection

**DOI:** 10.1101/2023.03.01.530674

**Authors:** Juan F. Quintana, Matthew C. Sinton, Praveena Chandrasegaran, Agatha Nabilla Lestari, Rhiannon Heslop, Bachar Cheaib, John Ogunsola, Dieudonne Mumba Ngoyi, Nono-Raymond Kuispond Swar, Anneli Cooper, Seth B. Coffelt, Annette MacLeod

**Affiliations:** Wellcome Centre for Integrative Parasitology (WCIP). University of Glasgow, Glasgow, UK. School of Biodiversity, One Health, Veterinary Medicine (SBOHVM), College of Medical, Veterinary and Life Sciences, University of Glasgow, Glasgow UK.; Lydia Becker Institute of Immunology and Inflammation, University of Manchester, UK.; Division of Immunology, Immunity to Infection and Health, Manchester Academic Health Science Centre, University of Manchester, UK.; Department of Parasitology, National Institute of Biomedical Research, Kinshasa, Democratic Republic of the Congo.; School of Cancer Sciences, University of Glasgow Glasgow, UK.; Cancer Research UK Beatson Institute, Glasgow, UK.

**Keywords:** African trypanosomes, γδ T cells, Vγ6^+^ cells, adipocytes, skin inflammation.

## Abstract

African trypanosome parasites colonise the skin in a process important for parasite transmission. However, how the skin responses to trypanosome infection remain unresolved. Here, using a combination of spatial and single cell transcriptomics, coupled with *in vivo* genetic models, we investigated the local immune response of the skin in a murine model of infection. First, we detected a significant expansion of IL-17A-producing γδ T cells (primarily Vγ6^+^) in the infected murine skin compared to naïve controls that occur mainly in the subcutaneous adipose tissue. Second, interstitial preadipocytes located in the subcutaneous adipose tissue upregulate several genes involved in inflammation and antigen presentation, including T cell activation and survival. *In silico* cell-cell communication suggests that adipocytes trigger γδ T cell activation locally *via Cd40, Il6, Il10,* and *Tnfsf18* signalling, amongst others. Third, mice deficient in IL-17A-producing γδ T cells show extensive inflammation, increased frequency of skin-resident IFNγ-producing CD8^+^ T cells and limited subcutaneous adipose tissue wasting compared to wild-type infected controls, independent of T_H_1 CD4^+^ T cells and parasite burden. Based on these observations, we proposed a model whereby adipocytes as well as Vγ6^+^ cells act concertedly in the subcutaneous adipose tissue to limit skin inflammation and tissue wasting. These studies shed light onto the mechanisms of γδ T cell-mediated immunity in the skin in the context of African trypanosome infection, as well as a potential role of immature and mature adipocytes as homeostatic regulators in the skin during chronic infection.

## Introduction

The skin represents the ultimate barrier, offering physical and mechanical protection against insults including infections. The skin is a complex organ encompassing several tissue layers, including the dermis and epidermis, which are capable of extensive remodelling and resistance to mechanical and chemical stimuli. Moreover, the skin hosts a complex stromal microenvironment containing a myriad of resident and recruited immune cells that actively survey the tissue. Amongst the dermal immune compartment, γδ T cells have emerged as critical regulators of tissue homeostasis^1–4^. For instance, Vγ6^+^ cells are enriched in mucosal tissues^3^ and the dermis^2, 4^ where they are considered resident cells. These skin resident γδ T cells express a wide range of effector molecules including interleukin 17A (IL-17A) and IL-17F^5^, IL-22^6^, and the epidermal growth factor (EGF) receptor ligand^7^, and amphiregulin (AREG)^8^, which are critical for promoting tissue repair and limiting inflammation following an insult^9–12^. Although γδ T cells have gathered a lot of attention for their role in cancer^13–16^, and autoimmune disorders affecting the skin such as psoriasis^17^ and lupus^18^, their function during parasitic colonisation of the skin is still not well understood.

Despite the robust nature of the skin as a barrier, some pathogens manage to circumvent its defences. Of these Trypanosomatids, including the causative agents of leishmaniasis, chagas disease, and sleeping sickness, are known to colonise the skin in a process that has been postulated as being critical for infection and parasite transmission^19–21^. Indeed, we and others have recently reported the presence of *Trypanosoma brucei* in the skin of mice and humans^21–23^. Such skin infections are often associated with dermatitis and pruritus^22^ or can occur in the absence of clinical symptoms or a positive result from gold-standard diagnostic methods^22^. This strongly indicates that mild or asymptomatic skin infections act as important but overlooked parasite reservoirs hampering ongoing eradication efforts. However, to date, the skin response to *T. brucei* infection remains unknown.

Here, we used a combined single cell and spatial transcriptomics approach to generate a spatially resolved single cell atlas of the murine skin during *T. brucei* infection. This combined approach led us to identify that interstitial preadipocytes and Langerhans cells both have central roles in the production of local cytokines and antigenic presentation, taking place largely in the subcutaneous adipose tissue and in proximity to the subcutaneous adipose tissue, respectively. Furthermore, we identified a population of skin IL-17A-producing Vγ6^+^ cells, located in proximity to adipocytes in the subcutaneous adipose tissue and subcutaneous adipose tissue, that display features of TCR engagement and T cell activation upon infection. Cell-cell communication analyses between subcutaneous adipocytes and Vγ6^+^ cells predicted several potential interactions mediating not only T cell activation *via Tnfsf18*, but also promoting adipocyte lipolysis *via* Vγ6^+^ cell-derived *Clcf1* and *Areg*. *In vivo* analyses revealed that Vγ4/6 γδ T cell-deficient mice experience severe skin inflammation, likely mediated by an increased frequency of IFNγ-producing CD8^+^ T cells in the skin, but with limited subcutaneous adipose tissue wasting compared to infected wild type controls. We conclude that IL-17A-producing Vγ6^+^ cells are critical mediators of skin immunity against African trypanosomes, dampening IFNγ-mediated CD8^+^ T cell responses in the skin and promoting subcutaneous adipose tissue wasting, supporting our recently identified role of IL-17 signalling as a mediator of adipose tissue wasting during *T. brucei* infection^24^. More broadly, our results provide a spatially resolved atlas that can help to dissect further immunological pathways triggered in the skin in response to infection.

## Materials and methods

### Ethical statement

All animal experiments were approved by the University of Glasgow Ethical Review Committee and performed in accordance with the home office guidelines, UK Animals (Scientific Procedures) Act, 1986 and EU directive 2010/63/EU. All experiments were conducted under SAPO regulations and UK Home Office project licence number PC8C3B25C to Dr. Jean Rodgers.

### Murine infections with *Trypanosoma brucei*

6–8-week-old wild-type female BALB/c (stock 000651) and FVB/NJ mice (stock 001800) were purchased from Jackson Laboratories. Vγ4/6^-/-^ mice (a gift from Rebecca O’Brien, National Jewish Health) were backcrossed to FVB/N. Female mice aged (6-8 weeks old) were used for infection. Six-to-eight-week-old wild-type female BALB/c (stock 000651) and FVB/NJ mice (stock 001800) were purchased from Jackson Laboratories. For infections, mice were inoculated by intra-peritoneal injection with ∼2 x 10^3^ parasites of strain *T. brucei brucei* Antat 1.1E^25^. **Click or tap here to enter text.**Parasitaemia was monitored by regular sampling from tail venepuncture and blood was examined using phase microscopy and the rapid “matching” method ^16^. Uninfected mice of the same strain, sex and age served as uninfected controls. Mice were fed *ad libitum* and kept on a 12 h light–dark cycle. All the experiments were conducted between 8h and 12h.

### Skin tissue processing and preparation of single cell suspension for single-cell RNA sequencing

Infected animals and naïve controls were anesthetized with isoflurane at 21 days post-infection and perfused transcardially with 25-30 ml of ice-cold 1X PBS containing 0.025% (wt/vol) EDTA. Skin biopsies from the abdominal flank area were shaved off completely. Some of these sections were placed in 4% PFA for 16 hours prior to embedding in paraffin and histological analysis. Single-cell dissociations for scRNAseq experiments were performed using the 2-step protocol published by (Joost *et al.*, 2020). Briefly, excised 4cm^2^ skin sections were diced into small pieces with a scalpel blade and enzymatically digested with Collagenase type I (500 U/ml; Gibco) and DNAse I (1 mg/ml; Sigma) in HBSS containing 0.04% BSA (Invitrogen) for ∼1 hr at 37 °C with shaking at 300rpm. Liberated cells from the partially digested tissue were pushed through a 70 μm nylon mesh filter with an equal volume of HBSS 0.04% BSA, and then kept on ice. The tissue remaining in the 70 μm filter was incubated in 0.05% trypsin EDTA for 15 minutes at 37°C, and then liberated cells pushed through the filter with an equal of HBSS 0.04% BSA. Flowthrough from the two digestion steps were combined, passed through a 40μM filter to remove any cell aggregate and spun at 350g for 10mins at 4°C. Finally, cells were passed through a MACS dead cell removal kit (Miltenyi Biotec) and diluted to ∼1,000 cells/μl in 200μl HBSS 0.04% BSA and kept on ice until single-cell capture using the 10X Chromium platform. The single cell suspensions were loaded onto independent single channels of a Chromium Controller (10X Genomics) single-cell platform. Briefly, ∼25,000 single cells were loaded for capture using 10X Chromium NextGEM Single cell 3 Reagent kit v3.1 (10X Genomics). Following capture and lysis, complementary DNA was synthesized and amplified (12 cycles) as per the manufacturer’s protocol (10X Genomics). The final library preparation was carried out as recommended by the manufacturer with a total of 14 cycles of amplification. The amplified cDNA was used as input to construct an Illumina sequencing library and sequenced on a Novaseq 6000 sequencers by Glasgow Polyomics.

### Read mapping, data processing, and integration

For FASTQ generation and alignments, Illumina basecall files (*.bcl) were converted to FASTQs using bcl2fastq. Gene counts were generated using Cellranger v.6.0.0 pipeline against a combined *Mus musculus* (mm10) and *Trypanosoma brucei* (TREU927) transcriptome reference. After alignment, reads were grouped based on barcode sequences and demultiplexed using the Unique Molecular Identifiers (UMIs). The mouse-specific digital expression matrices (DEMs) from all six samples were processed using the R (v4.2.1) package Seurat v4.1.0^17^. Additional packages used for scRNAseq analysis included dplyr v1.0.7 ^18^, RColorBrewer v1.1.2 (http://colorbrewer.org), ggplot v3.3.5 ^19^, and sctransform v0.3.3 ^20^. We initially captured 65,734 cells mapping specifically against the *M. musculus* genome across all conditions and biological replicates, with an average of 29,651 reads/cell and a median of 1,762 genes/cell (**S1A Table and Supplementary Figure 1**). The number of UMIs was then counted for each gene in each cell to generate the digital expression matrix (DEM). Low quality cells were identified according to the following criteria and filtered out: *i*) nFeature > 100 or <5,000, *ii*) nCounts > 100 or <20,000, *iii*) > 30% reads mapping to mitochondrial genes, and *iv*) > 40% reads mapping to ribosomal genes, v) genes detected < 3 cells. After applying this cut-off, we obtained a total of 56,876 high quality mouse-specific cells with an average of 29,651 reads/cells and a median of 1,683 genes/cell (**S1A Table**). High-quality cells were then normalised using the *SCTransform* function, regressing out for total UMI and genes counts, cell cycle genes, and highly variable genes identified by both Seurat and Scater packages, followed by data integration using *IntegrateData* and *FindIntegrationAnchors*. For this, the number of principal components were chosen using the elbow point in a plot ranking principal components and the percentage of variance explained (15 dimensions) using a total of 5,000 genes, and SCT as normalisation method.

### Cluster analysis, marker gene identification, and subclustering

The integrated dataset was then analysed using *RunUMAP* (10 dimensions), followed by *FindNeighbors* (10 dimensions, reduction = “pca”) and *FindClusters* (resolution = 0.4). With this approach, we identified a total of 16 cell clusters The cluster markers were then found using the *FindAllMarkers* function (logfc.threshold = 0.25, assay = “RNA”). To identify cell identity confidently, we employed a supervised approach. This required the manual inspection of the marker gene list followed by and assignment of cell identity based on the expression of putative marker genes expressed in the unidentified clusters. The resolution used for these analyses was selected using the Clustree package^26^ **(Supplementary Figure 1).** A cluster name denoted by a single marker gene indicates that the chosen candidate gene is selectively and robustly expressed by a single cell cluster and is sufficient to define that cluster (e.g., *Col1a1, Cd3g, Pparg, Krt14*, among others). When manually inspecting the gene markers for the final cell types identified in our dataset, we noted the co-occurrence of genes that could discriminate two or more cell types (e.g., T cells, fibroblasts). To increase the resolution of our clusters to help resolve potential mixed cell populations embedded within a single cluster, we subset fibroblasts, myeloid cells, and T cells and analysed them separately using the same functions described above. In all cases, upon subsetting, the resulting objects were reprocessed using the functions *FindVariableFeatures, ScaleData, RunUMAP, FindNeighbors*, and *FindClusters* with default parameters. The number of dimensions used in each case varied depending on the cell type being analysed but ranged between 5 and 10 dimensions. Cell type-level differential expression analysis between experimental conditions was conducted using the *FindMarkers* function (*min.pct* = 0.25, *test.use* = Wilcox) and (*DefaultAssay* = “SCT”). Cell-cell interaction analysis mediated by ligand-receptor expression level was conducted using NicheNet ^25^ with default parameters using “mouse” as a reference organism, comparing differentially expressed genes between experimental conditions (*condition_oi* = “Infected”, *condition_reference* = “Uninfected”).

### 10X Visium spatial sequencing library preparation and analysis

#### Tissue processing and library preparation

Mice were shaved and dissected skin stored in 4% paraformaldehyde for 24 h, before paraffin embedding. RNA was purified from FFPE sections to measure integrity, using the Absolutely Total RNA FFPE Purification Kit (Agilent Technologies) as per the manufacturer’s instructions. RNA integrity was then measured by BioAnalyzer (Agilent Technologies) using the RNA 6000 Pico Kit (Agilent Technologies). Samples selected for sequencing all had a DV200 >50%. We then placed 10 µm sections within the capture areas of Visium Spatial Slides (10X Genomics) and proceeded to perform H&E staining and image capture, as per the manufacturer’s instructions. To deparaffinise, slides were incubated at 60 °C for 2 h before incubating twice in xylene at room temperature, for 10 min each, three times in 100% ethanol for 3 min each, twice in 96% ethanol for 3 min each, 85% ethanol for 3 min, and then submerged in Milli-Q water until ready to stain. For H&E staining, slides were incubated with Mayer’s haematoxylin Solution (Sigma-Aldrich), Bluing Buffer (Dako) and Alcoholic Eosin solution (Sigma-Aldrich), with thorough washing in ultrapure water between each step. Stained slides were scanned under a microscope (EVOS M5000, Thermo). De-crosslinking was performed by incubating sections twice with 0.1 N HCl for 1 min at room temperature. Sections were then incubated twice with Tris-EDTA (TE) buffer pH 9.0 before incubating at 70 °C for 1 h. The Visium Spatial Gene Expression Slide Kit (10X Genomics) was used for reverse transcription and second strand synthesis, followed by denaturation, to allow the transfer of the cDNA from the slide to a collection tube. These cDNA fragments were then used to construct spatially barcoded Illumina-compatible libraries using the Dual Index Kit TS Set A (10x Genomics) was used to add unique i7 and i5 sample indexes, enabling the spatial and UMI barcoding. The final Illumina-compatible sequencing library underwent paired-end sequencing (150 bp) on a NovaSeq 6000 (Illumina) at GenomeScan.

After sequencing, the FASTQ files were aligned to a merged reference transcriptome combining the Mus musculus genome (mm10). After alignment, reads were grouped based on spatial barcode sequences and demultiplexed using the UMIs, using the SpaceRanger pipeline version 1.2.2 (10X Genomics). Downstream analyses of the expression matrices were conducted using the Seurat pipeline for spatial RNA integration (Hao et al., 2021b; Stuart et al., 2019) (Table S3A). Specifically, the data was scaled using the SCTransform function with default parameters. We then proceeded with dimensionality reduction and clustering analysis using *RunPCA* (assay = “SCT”), *FindNeighbours* and *FindClusters* functions with default settings and a total of 30 dimensions. We then applied the *FindSpatiallyVariables* function to identify spatially variable genes, using the top 1,000 most variable genes and “markvariogram” as selection method. To integrate our skin scRNAseq with the 10X Visium dataset, we used the *FindTransferAnchors* function with default parameters, using SCT as normalization method. Then, the *TransferData* function (weight.reduction = “pca”, 30 dimensions) was used to annotate brain regions based on transferred anchors from the scRNAseq reference datasets. To predict the cell-cell communication mediated by ligand-receptor co-expression patterns in the spatial context, we employed NICHES v0.0.2 (Raredon et al., 2022). Upon dimensionality reduction and data normalisation, NICHES was run using fanton5 as ligand-receptor database with default parameters. The resulting object was then scaled using the functions *ScaleData*, *FindVariableFeatures* (selection.method = “disp”), RunUMAP with default settings and a total of 15 dimensions. Spatially resolved expression of ligand-receptor pairs was then identified using the *FindAllMarkers* function (min.pct = 0.25, test.use = “roc”). For visualisation, we used the *SpatialFeaturePlot* function with default parameters and min.cutoff = “q1”.

#### Single molecule fluorescence *in situ* hybridisation (smFISH) using RNAscope

smFISH experiments were conducted as follows. Briefly, to prepare tissue sections for smFISH, infected animals and naïve controls were anesthetized with isoflurane, and skin sections from the flank were shaved off completely before placing in 4% PFA overnight prior to embedding in paraffin. Paraffin-embedded skin sections sections (3 mm) were prepared placed on a SuperFrost Plus microscope slides. Sections were then then dehydrated in 50, 70 and 100% ethanol. RNAscope 2.5 Assay (Advanced Cell Diagnostics) was used for all smFISH experiments according to the manufacturer’s protocols. All RNAscope smFISH probes were designed and validated by Advanced Cell Diagnostics. For image acquisition, 16-bit laser scanning confocal images were acquired with a 63x/1.4 plan-apochromat objective using an LSM 710 confocal microscope fitted with a 32-channel spectral detector (Carl Zeiss). Lasers of 405nm, 488nm and 633 nm excited all fluorophores simultaneously with corresponding beam splitters of 405nm and 488/561/633nm in the light path. 9.7nm binned images with a pixel size of 0.07um x 0.07um were captured using the 32-channel spectral array in Lambda mode. Single fluorophore reference images were acquired for each fluorophore and the reference spectra were employed to unmix the multiplex images using the Zeiss online fingerprinting mode. smFISH images were acquired with minor contrast adjustments as needed, and converted to grayscale, to maintain image consistency.

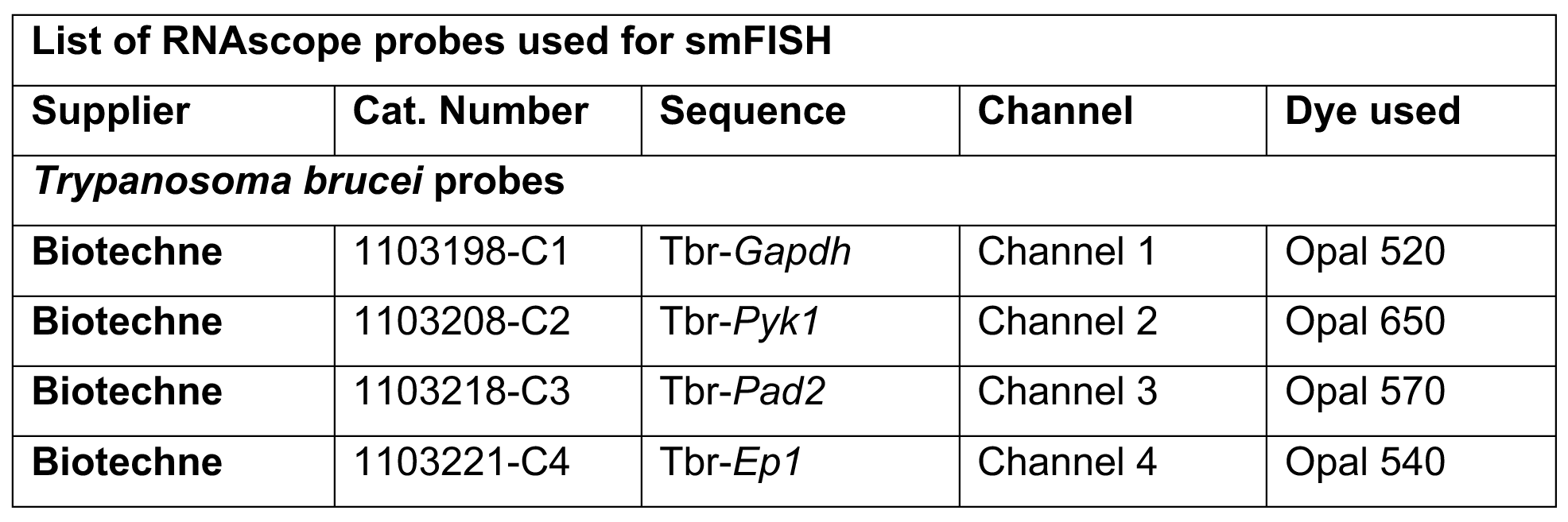

| List of RNAscope probes used for smFISH |  |  |  |  |
| --- | --- | --- | --- | --- |
| Supplier | Cat. Number | Sequence | Channel | Dye used |
| ***Trypanosoma brucei* probes** |  |  |  |  |
| Biotechne | 1103198-C1 | Tbr-*Gapdh* | Channel 1 | Opal 520 |
| Biotechne | 1103208-C2 | Tbr-*Pyk1* | Channel 2 | Opal 650 |
| Biotechne | 1103218-C3 | Tbr-*Pad2* | Channel 3 | Opal 570 |
| Biotechne | 1103221-C4 | Tbr-*Ep1* | Channel 4 | Opal 540 |

#### Histological analysis of adipocyte size

Skin placed into 4% paraformaldehyde (PFA) and fixed overnight at room temperature. PFA-fixed skin biopsies were then embedded in paraffin, sectioned, and stained by the Veterinary Diagnostic Services facility (University of Glasgow, UK). Sections were Haematoxylin and Eosin (H&E) stained for adipocyte size analysis. Slide imaging was performed by the Veterinary Diagnostic Services facility (University of Glasgow, UK) using an EasyScan Infinity slide scanner (Motic, Hong Kong) at 20X magnification. To determine subcutaneous adipocyte size in skin sections, images were first opened in QuPath (v. 0.3.2)^27^, before selecting regions and exporting to Fiji^28^. In Fiji, images were converted to 16-bit format, and we used the Adiposoft plugin^29^ to quantify adipocyte size within different sections.

#### Semi-quantitative evaluation of the parasite burden and inflammation in skin sections

Paraffin-embedded skin samples were cut into 2.5 μm sections and stained for *T*. *brucei* parasites using a polyclonal rabbit antibody raised against *T*. *brucei* luminal binding protein 1 (BiP) (J. Bangs, SUNY, USA) using a Dako Autostainer Link 48 (Dako, Denmark) and were subsequently counterstained with Gill’s Haematoxylin. The extent of inflammatory cell infiltration in skin sections was assessed in haematoxylin and eosin-stained sections. Stained slides were assessed by two independent pathologists blinded to infection status and experimental procedures. Skin parasite burden was assessed for both intravascular (parasites within the lumen of dermal or subcutaneous small to medium-sized vessels) and extravascular locations (parasites outside blood vessels, scattered in the connective tissue of the dermis or in the subcutaneous adipose tissue) and was evaluated in 5 randomly selected fields at 40X magnification for each sample. A semi-quantitative ordinal score was used to grade the trypanosomes burden in the skin, as follows: 0 = no parasites; 1 = 1 to 19 trypanosomes; 2 = 20-50 trypanosomes; 3 = > 50 trypanosomes. An average parasite burden score per field of view was calculated for each skin section. The severity of skin inflammation was assessed semi-quantitatively, graded on a 0–3 scale: 0 (leukocytes absent or rarely present); 1, mild (low numbers of mixed inflammatory cells present); 2, moderate (increased numbers of mixed inflammatory cells), and 3, marked (extensive numbers/aggregates of inflammatory cells). The average of 10 randomly selected fields at 20X objective magnification for each skin section determined the inflammatory score.

#### DNA extraction and Quantitative Polymerase Chain Reaction (qPCR) of *T. brucei* in murine skin tissue

Genomic DNA (gDNA) was extracted from 25–30 mg skin tissue preserved at −80°C. After defrosting on ice, skin was finely chopped with scissors, and disrupted for 8 minutes in 300 μL ATL buffer (Qiagen) using a Qiagen Tissuelyser at 50Hz with a ceramic bead (MPBio). Disrupted tissues was incubated at 56°C with 2mg/ml proteinase K (Invitrogen) overnight and DNA extracted from digested tissue using the Qiagen DNeasy Blood and Tissue Kit (Qiagen, Manchester (UK). The resulting gDNA was quantified using a Qubit Fluorimeter (Thermofisher Scientific). and diluted to 4 ng/μl. Trypanosome load in the skin was determined using Taqman real-time PCR, using primers and probe specifically designed to detect the trypanosome *Pfr2* gene, as previously reported^30^. Reactions were performed in a 25 μl reaction mix comprising 1X Taqman Brilliant III master mix (Agilent, Stockport, UK), 0.2 pmol/μl forward primer (CCAACCGTGTGTTTCCTCCT), 0.2 pmol/μl reverse primer (CGGCAGTAGTTTGACACCTTTTC), 0.1 pmol/μl probe (FAM-CTTGTCTTCTCCTTTTTTGTCTCTTTCCCCCT-TAMRA) (Eurofins Genomics, Germany) and 20 ng template DNA. A standard curve was constructed using a serial 10-fold dilution range: 1x 10^6^ to 1x 10^1^ copies of PCR 2.1 vector containing the cloned *Pfr2* target sequence (Eurofins Genomics, Germany). The amplification was performed on an ARIAMx system (Agilent, USA) with a thermal profile of 95°C for 3 minutes followed by 45 cycles of 95°C for 5 seconds and 60°C for 10 seconds. The *Pfr2* copy number within each 20ng DNA skin sample was calculated from the standard curve using the ARIAMx qPCR software (Agilent, USA) as a proxy for estimated trypanosome load. Skin biopsies from naïve controls were included to determine the background signal and detection threshold.

#### Flow cytometry analysis of skin-dwelling T lymphocytes

Murine skin biopsies were harvested and digested as indicated above. The resulting single cell suspensions were resuspended in ice-cold FACS buffer (2 mM EDTA, 5 U/ml DNAse I, 25 mM HEPES and 2.5% Foetal calf serum (FCS) in 1× PBS), blocked with TruStain FcX (anti-mouse CD16/32) antibody (Biolegend, 1:1,000), and stained for extracellular markers at 1:400 dilution. Macrophages (F4/80), B cells (CD19), and erythrocytes (TER119) were excluded from the analysis by included them in a dump channel. For intracellular staining, single-cell suspensions were stimulated as above in Iscove’s modified Dulbecco’s media (supplemented with 1× non-essential amino acids, 50 U/ml penicillin, 50 μg/ml streptomycin, 50 μM β-mercaptoethanol, 1 mM sodium pyruvate and 10% FBS. Gibco) containing 1X cell Stimulation cocktail containing phorbol 12-myristate 13-acetate (PMA), Ionomycin, and Brefeldin A (eBioSciences^TM^) for 5 hours at 37°C and 5% CO2. Unstimulated controls were also included in these analyses. After surface marker staining, cells were fixed and permeabilized with a Foxp3/Transcription Factor Staining Buffer Set (eBioscience) and stained overnight at 4 °C. The list of flow cytometry antibodies used in this study were obtained from Biolegend and are presented in the table below. For flow cytometry analysis, samples were run on a flow cytometer LSRFortessa (BD Biosciences) and analysed using FlowJo software version 10 (Treestar).

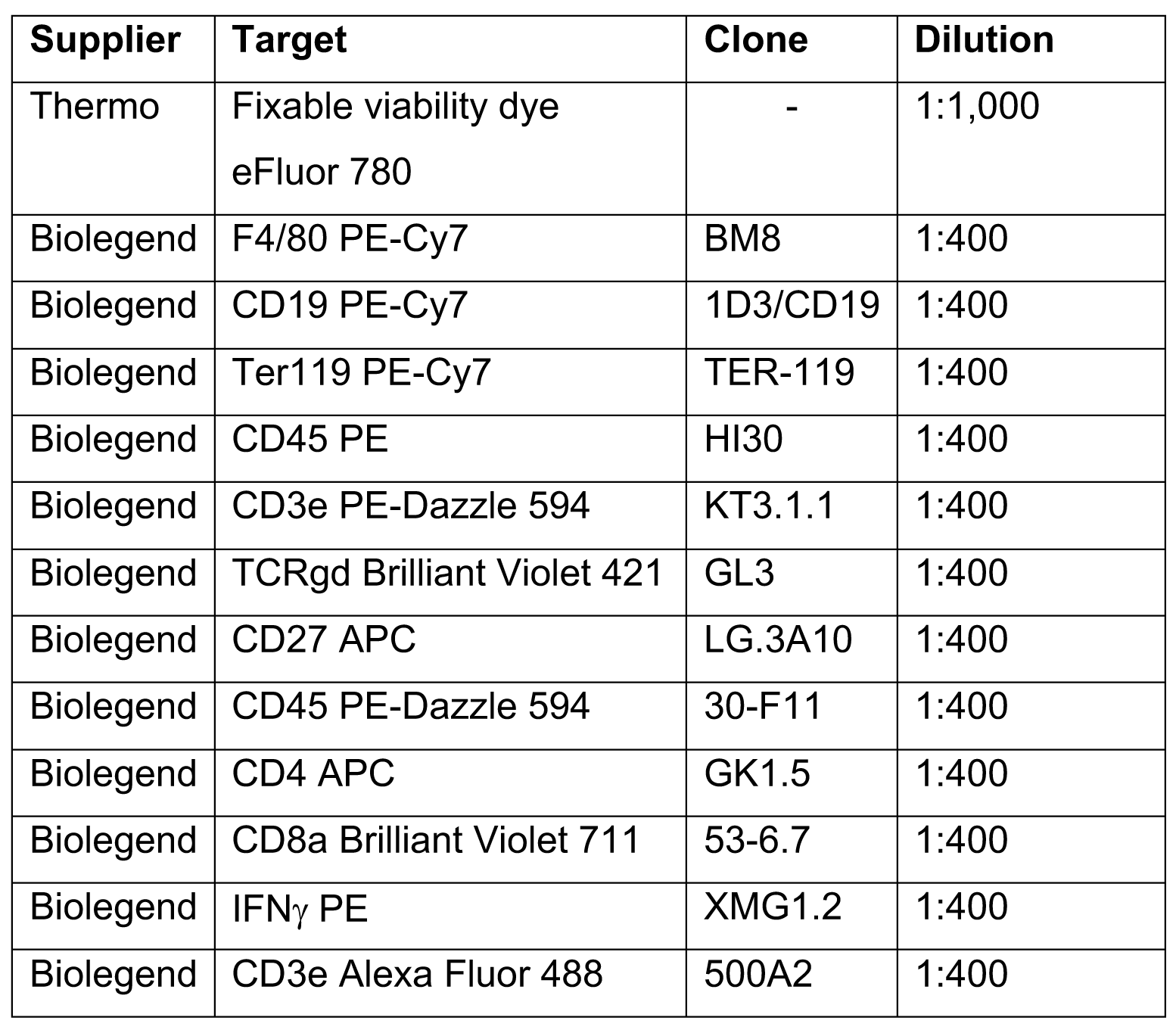

| Supplier | Target | Clone | Dilution |
| --- | --- | --- | --- |
| Thermo | Fixable viability dye<br>eFluor 780 | - | 1:1,000 |
| Biolegend | F4/80 PE-Cy7 | BM8 | 1:400 |
| Biolegend | CD19 PE-Cy7 | 1D3/CD19 | 1:400 |
| Biolegend | Ter119 PE-Cy7 | TER-119 | 1:400 |
| Biolegend | CD45 PE | HI30 | 1:400 |
| Biolegend | CD3e PE-Dazzle 594 | KT3.1.1 | 1:400 |
| Biolegend | TCRgd Brilliant Violet 421 | GL3 | 1:400 |
| Biolegend | CD27 APC | LG.3A10 | 1:400 |
| Biolegend | CD45 PE-Dazzle 594 | 30-F11 | 1:400 |
| Biolegend | CD4 APC | GK1.5 | 1:400 |
| Biolegend | CD8a Brilliant Violet 711 | 53-6.7 | 1:400 |
| Biolegend | IFNγ PE | XMG1.2 | 1:400 |
| Biolegend | CD3e Alexa Fluor 488 | 500A2 | 1:400 |

## Data availability

The transcriptome data generated in this study have been deposited in the Gene Expression Omnibus (GSE226113). The processed transcript count data and cell metadata generated in this study are available at Zenodo (DOI: 10.5281/zenodo.7677469). Additional data and files can also be sourced via Supplementary Tables.

## Code availability

Code used to perform analysis described can be accessed at Zenodo (DOI: 10.5281/zenodo.7677469).

## Results

### Spatially resolved single cell atlas of the murine skin chronically infected with African trypanosomes

To study the immune responses in the murine skin against African trypanosomes, we conducted a combined single cell and spatial transcriptomic analysis using 10X Genomics (**Materials and methods**). Skin from female BALB/c mice infected with *T. brucei* Antat 1.1 displayed marked inflammation, as determined by H&E examination (**Figure 1A**), compared to uninfected skin. Parasites were detected in extravascular spaces (**Figure 1B**) at 25 days post-infection, as previously reported^21^. We chose to explore this time point as the *T. brucei* skin infection is well established, enabling us to explore how the skin adapts to such conditions. Following skin dissociation and scRNAseq analysis, we obtained a total of 56,876 high-quality cells with an average of 1,622 genes and 29,651 reads per cell (**Figure 1C, S1 Figure, and S1 Table**) from naïve (*n* = 2) and infected (*n* = 2) mice. These cells were broadly classified into 16 different clusters (**Figure 1C**) based on common expression makers putatively associated with these clusters (**Figure 1D**). The stromal compartment consisted of eight keratinocyte clusters (46,591 cells; KC 1 to 8), characterised by a high expression level for *Krt14, Krt15, Krt35, Krt25, and Fabp5*, in addition to one cluster with high expression of *Col1a1, Ly6a, Thy1, Pdgfra*, and *Cd34* that we designated as fibroblast-like mesenchymal cells (4,274 cells), one *Cd36^+^ Cldn5^+^ Pecam1^+^* endothelial cell cluster (792 cells), one *Pparg^+^* adipocyte (529 cells), and one *Mlana^+^* melanocyte cluster (755 cells) (**Figure 1C and D**). In the spatial context, the keratinocyte populations were mainly detected in the dermis and epidermis, whereas the rest of the stromal cells were detected in the muscle layer and subcutaneous adipose tissue in both naïve and infected samples (**Figure 1E, S2 Figure, and S2 Table**).

**Figure 1.**
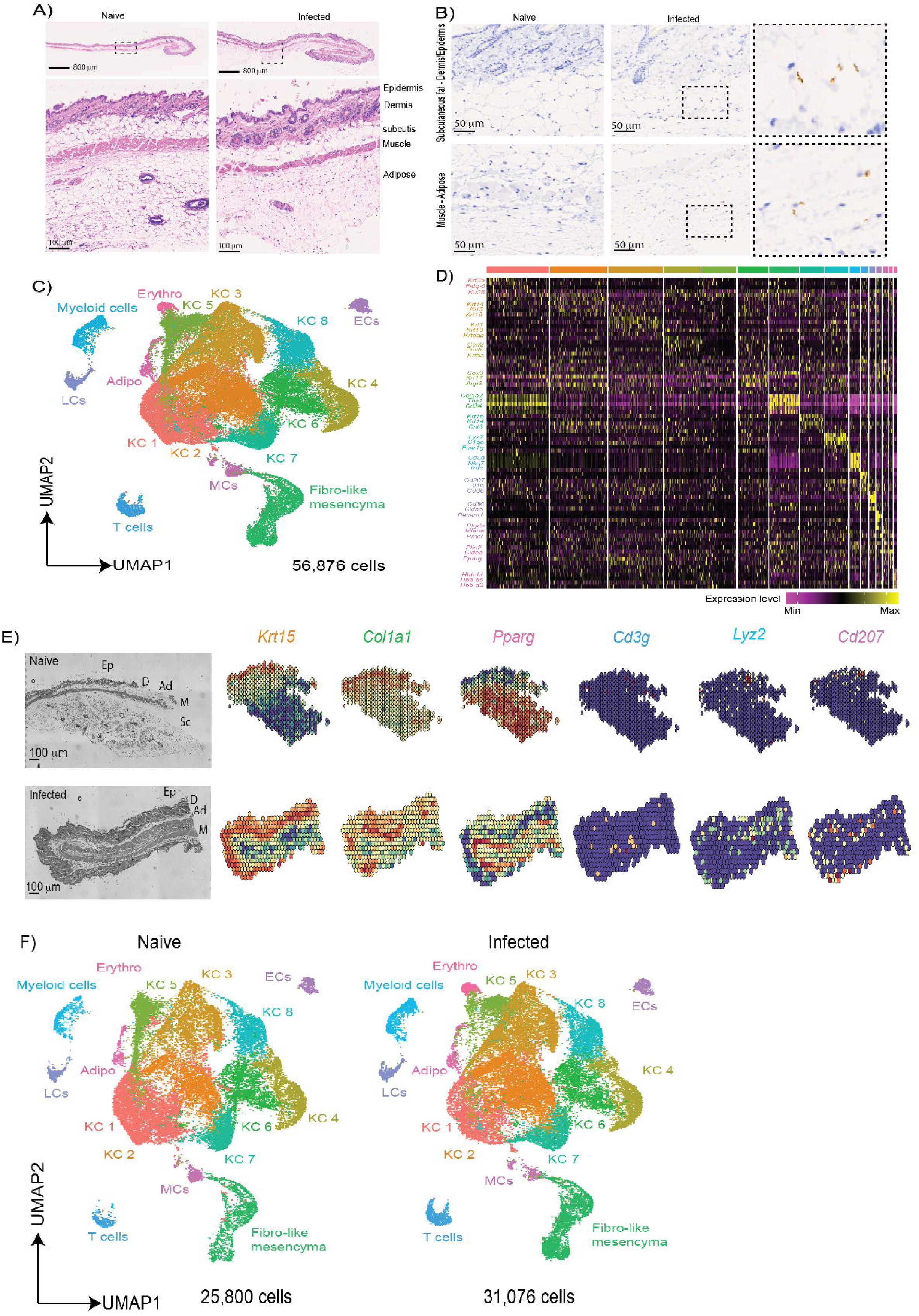
Integrative overview of the murine skin infected with *T. brucei* infection using single cell transcriptomics. **A)** H&E-stained images from naïve and infected mice. Scale bar: 800 μm (top) and 100 μm (bottom). **B)** Immunohistochemistry against the stumpy-specific marker PAD1 in naïve and infected samples, including insets highlighting the presence of stumpy forms in both the epidermis-subcutaneous adipose tissue and the subcutaneous adipose tissue-adipose tissue. Scale bar: 50 μm. **C)** Uniform manifold approximation and projection (UMAP) of 56,876 high-quality cells from both naïve and infected samples, highlighting the major cell types detected in our dataset, including stromal cells (Keratinocytes, fibroblasts, endothelial cells, and adipocytes) and immune cells (myeloid cells, T cells, Langerhans cells, and macrophages). **D)** Top 10 marker genes defining all the major cell clusters detected in (C). The heatmap is colour-coded based on gene expression intensity. **E)** Spatial feature plot for selected marker genes defining keratinocytes (*Krt15*), fibroblast-like mesenchymal cells (*Col1a1*), adipocytes (*Pparg*), T cells (*Cd3g*), myeloid cells (*Lyz2*), and Langerhans cells (*Cd207*). The corresponding histological section is included on the left, including an annotation of epidermis (Ep), dermis (D), adipose tissue (Ad), Muscle (M), and subcutaneous adipose tissue (Sc). **F)** As in (C) but splitting the UMAP plot into naïve samples (25,800 cells; *n* = 2 mice) and infected (31,076 cells; *n* = 2). KC: keratinocytes, FB: Fibroblasts, EC: endothelial cells, MCs: Macrophages, LC: Langerhans cells, Adipo: Adipocytes, Erythro: Erythrocytes.

The immune compartment consisted of *Cd3g^+^ Trdc^+^* T cells (1,103 cells), *Cd207^+^* Langerhans cells (854 cells), *Lyz2^+^* myeloid cells (1,499 cells), and erythrocytes (479 cells) (**Figure 1C and 1D**). The majority of these immune cells were lowly detected in naïve skin but were readily found in the subcutaneous adipose tissue of infected samples (**Figure 1E**). A closer examination of differences between experimental conditions allowed us to detect the expansion of individual cell clusters in response to infection. For example, the *Lyz2^+^* myeloid and *Cd3g^+^* T cell clusters have higher frequencies in the infected skin compared to naïve controls (**Figure 1F**). Similarly, we noted the presence of a small erythrocyte cluster exclusively in the infected sample, which may be indicative of infection-induced vascular leakage (**Figure 1F**). These results are consistent with the increased frequency of inflammatory cells in infected samples that we determined by histopathological examination.

### Skin preadipocytes and keratinocytes provide inflammatory signals and antigen presentation during chronic *T. brucei* infection

Stromal cells are increasingly recognised as critical for initiating immunological responses in many tissues, including the skin^31^. To capture as much stromal cell resolution as possible, and to better understand how this compartment in the murine skin responds to infection, we reclustered stromal cell clusters (keratinocytes, adipocytes, endothelial cells, melanocytes, and fibroblast-like mesenchymal cells) and reanalysed them separately. After re-clustering, we obtained a total of 13 different stromal clusters, seven of which were *bona fide* keratinocytes expressing *Krt25, Krt14*, and *Krt15* (**Figure 2A and 2B**). In addition to these clusters, we also detected *Mlana^+^* melanocytes, *Plin2^+^* mature adipocytes, and *Cldn5^+^* endothelial cells (**Figure 2A and 2B**). Interestingly, the re-clustering analysis led us to identify a clear population of cells expressing bona fide mesenchymal genes including *Cd34, Thy1, Ly6a, Pdgfra, Pdgfrb,* and *Col1a1*, which we previously assigned as a fibroblast population (**Figure 2A and 2B**). We assigned these cells as interstitial preadipocytes (stem-like adipocyte precursor cells) as they also express *Dpp4*, *Pi16*, and *Dcn* (**Figure 2A and B**), markers previously shown to be upregulated in this cell type^32^. Some stromal populations were altered during infection. For instance, the frequency of KC2 and KC3 increased from around 15.02% and 10.92% to approximately 19.3% and 14.07%, respectively. In contrast, the adipocyte population decreased from 1.60% to 1.21%, indicating subcutaneous adipose tissue wasting (**Figure 2C**), as previously reported^24, 33^. We also observed a higher frequency of endothelial cells (ECs; from 6.67% to 8.02%) and interstitial preadipocyte population 2 (IPA2; from 1.98% to 3.81%) in infected compared with naïve skin (**Figure 2C**). Together this indicates that during *T. brucei* infection there is a remodelling of the stromal compartment within the skin. Stromal cells are recognised as drivers of inflammatory responses via the production of cytokines and chemokines^31^, as well as the induction of T cell activation *via* antigen presentation^34^. To unbiasedly assess our transcriptomics dataset, to determine whether stromal cells were influencing immune function, we performed module scoring. This was used to determine the global expression levels of cytokines (**Figure 2D**) and antigen presentation molecules (**Figure 2E**). This analysis revealed that IPA2 (interstitial preadipocyte 2), and to a lesser extent KC2 (keratinocyte 2), were the main producers of cytokines and antigen presentation molecules (**Figure 2D and 2E**). Indeed, almost exclusively in IPA2, we noted increased expression of *Il6, Il15, Il18bp* and interferon-driven *Cxcl9, Cxcl10*, as well as the class Ib and II major histocompatibility complex *H2-M3* and *H2-DMa*, respectively (**Figure 2F-2H**). In the spatial context, most of these responses were identified as restricted foci in the dermis and epidermis of naïve mice (**Figure 2I**), but significantly localised to the subcutaneous adipose tissue in infected mice (**Figure 2I**). Taken together, these results indicate that the IPA2 subcluster, located in the subcutaneous adipose tissue, is a key driver of recruitment and activation of immune cells in the skin in response to *T. brucei* infection.

**Figure 2.**
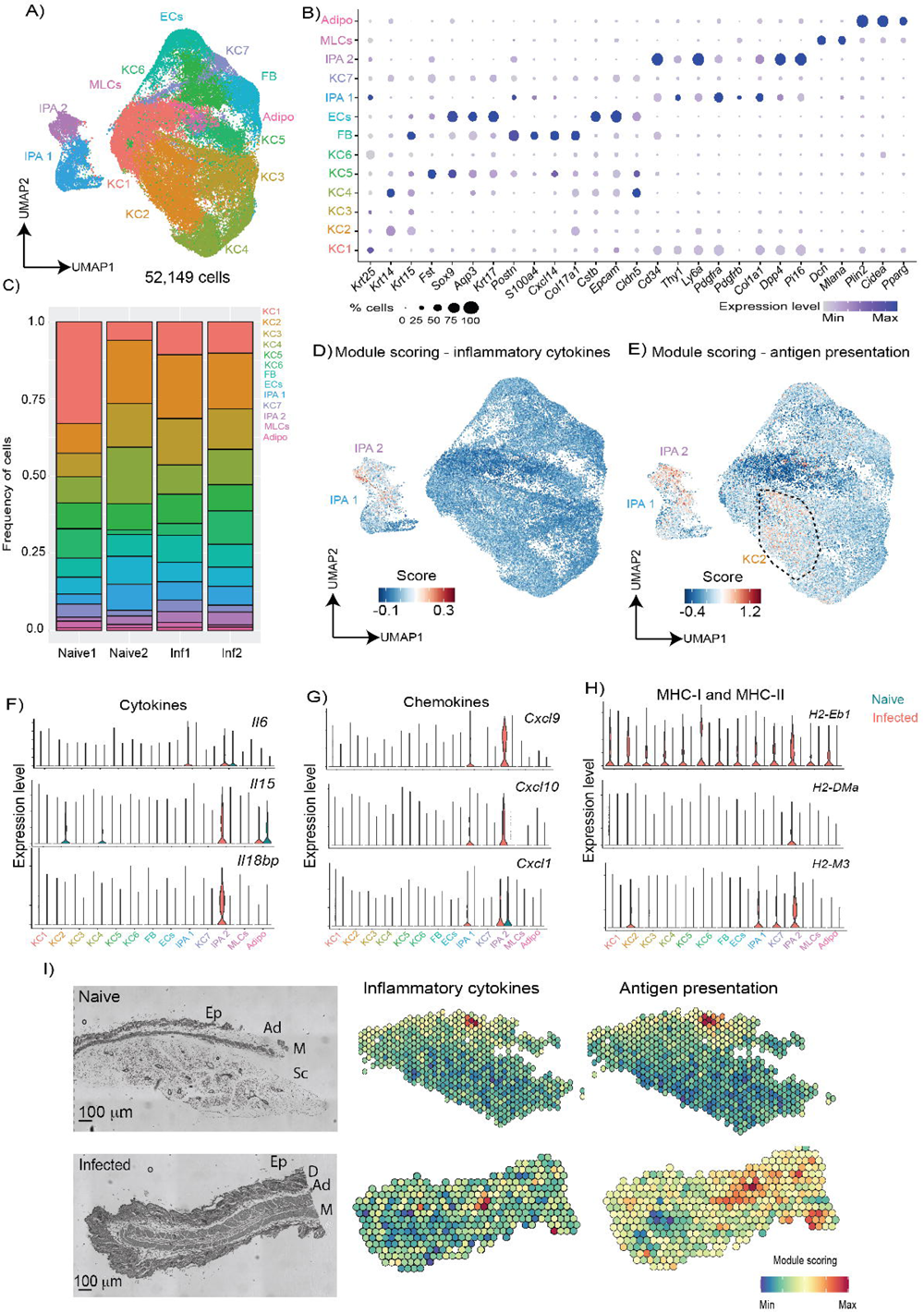
The murine skin stromal cells respond to infection by upregulating genes associated with antigen presentation and chemotaxis. **A)** Uniform manifold approximation and projection (UMAP) of 52,149 high-quality cells within the stromal subcluster, encompassing seven keratinocyte clusters (KC1-7), fibroblasts (FB), endothelial cells (ECs), melanocytes (MLCs), and interstitial preadipocytes (IPAs). **B)** Dot plot representing the expression levels of top marker genes used to catalogue the diversity of skin stromal cells. The side of the dots represent the percentage of cells that express a given marker, and the colour intensity represent the level of expression. **C)** Frequency plot the different stromal cell types detected in the murine skin in naïve (*n* = 2) and infected (*n* = 2) samples. Module scoring for the overall expression of inflammatory cytokines **(D)** and genes associated with antigen presentation **(E).** Violin plot showing the expression level of several significant adipocyte-specific cytokines **(F)**, chemokines **(G)**, and major histocompatibility complex (MHC) I and II molecules **(H). I)** Spatial module scoring for inflammation and antigen presentation in a naïve (top) and infected (bottom) skin section. Scale bar, 100 μm.

### Langerhans cells, dendritic cells, and *Cd14^+^* monocytes contribute to antigen presentation in the murine skin during chronic *T. brucei* infection

In addition to the stroma, resident myeloid cells are also involved in the initial responses to infection. Thus, we next examined transcriptional responses within the myeloid compartment, encompassing *Lyz2^+^* myeloid and *Cd207^+^* Langerhans cells (**Figure 1C and 1D**). To gain as much granularity as possible within this compartment, we reclustered the myeloid subset leading to the identification of six subclusters including *Cd14^+^* monocytes, *Mrc1^+^* macrophages, two clusters of *Cd207^+^* Langerhans cells, *Hdc*^+^ *Gata2*^+^ mast cells, and *Clec9a*^+^ *Btla*^+^ *Xcr1*^+^ dendritic cells, which we classed as cDC1s (**Figure 3A and 3B**). Interestingly, *Mrc1^+^* macrophages also expressed *Il10*, suggesting that these cells may have anti-inflammatory properties (**Figure 3B**). We found that upon infection, there is an expansion of all the myeloid cells, but notably so within the *Mrc1^+^* macrophage and *Cd14^+^* monocyte and Langerhans cell subclusters (**Figure 3A**). These subsets, as well as the mast cells and the cDC1s localise to the subcutaneous adipose tissue (**Figure 3C**), mirroring the localisation of other stromal cells driving immune cell recruitment and activation (**Figure 2I**). Given their role in initiating an adaptive immune response, we next focussed on characterising myeloid cell function by measuring the global expression levels of pro-inflammatory cytokines and molecules associated with antigen presentation, as we did for the stromal compartment, using module scoring. This analysis revealed that the *Cd14^+^* monocytes and *Mrc1*^+^ macrophages express the highest levels of cytokines within the myeloid compartment (**Figure 3D**). In contrast, antigenic presentation was predominantly driven by Langerhans cells and DCs (**Figure 3E**). Taken together, these analyses reveal that myeloid cells located in the subcutaneous adipose tissue drive the expression of pro-inflammatory cytokines and antigen presentation concertedly with stromal cells during infection.

**Figure 3.**
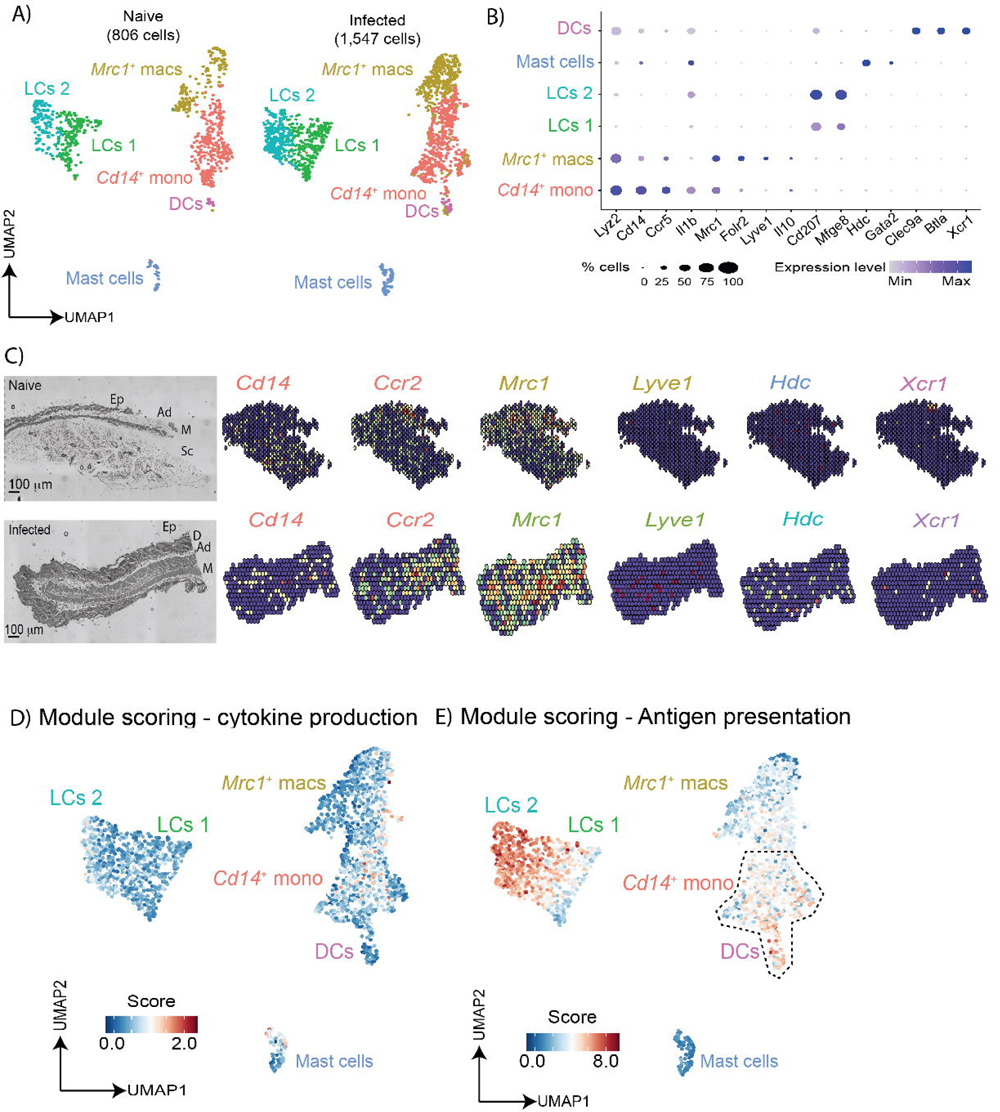
The murine skin is colonised by a myriad of myeloid cells during chronic *T. brucei* infection. **A)** Uniform manifold approximation and projection (UMAP) of 2,353 high-quality cells within the myeloid cluster were re-analysed to identify a total four subclusters, including dendritic cells (DCs), Mast cells, and two populations of macrophages (*Mrc1^+^* macs and *Cd14^+^* mono), and two populations of Langerhans cells (LCs 1 and LCs 2). **B)** Dot plot representing the expression levels of top marker genes used to catalogue the diversity of myeloid cells. **C)** Spatial feature plot depicting the expression levels of markers defining *Cd14^+^* monocytes (*Cd14, Ccr2*), *Mrc1^+^* macrophages (*Mrc1, Lyve1*), mast cells (*Hdc*), and DCs (*Xcr1*). The corresponding histological section is included on the left, including an annotation of epidermis (Ep), dermis (D), adipose tissue (Ad), Muscle (M), and subcutaneous adipose tissue (Sc). Module scoring for the overall expression of inflammatory cytokines **(D)** and genes associated with antigen presentation **(E)**.

### Both *Cd4*^+^ T cells and Vγ6^+^ cells expand in the skin during chronic *T. brucei* infection

We next examined the T cell compartment in our scRNAseq dataset. After re-clustering, we identified a total of 1,043 cells encompassing *Il5^+^ Gata3^+^* ILC2s (138 cells), *Ncr1^+^* NK cells (172 cells), *Icos^+^ Rora^+^* CD4*^+^* T cells (426 cells), and two separate cell clusters of *Trdc^+^* γδT cells, 1 (155 cells) and 2 (152 cells) (**Figure 4A and 4B**). Both of the γδ T cell clusters express high levels of *Tcrg-C1, Cd163l1*, but low levels of *Cd27* (**Figure 4B**), and based on recent literature, this indicates that they are likely to be IL-17-producing Vγ6^+^ cells^35^. *In vivo*, we observed a significant expansion of CD27^-^ (IL-17A-producing) γδ T cells and a concomitant reduction in the frequency of CD27^+^ (IFNγ-producing) γδ T cells in skin biopsies from infected BALB/c mice compared to naïve controls (**Figure 4C**), following the same trend predicted by the scRNAseq data (**Figure 4A and 4B**), thus validating our *in silico* prediction. Intriguingly, we noted that the Vγ6^+^ cell cluster 1 was only present in the naïve skin but disappeared upon infection (**Figure 4A**). In contrast, the Vγ6^+^ cell cluster 2 was not present in the naïve skin but appeared during infection (**Figure 4A**). Based on this finding, we hypothesised that these two γδT cell clusters represent different activation states, whereby γδT cells within cluster 1 represent “resting” cells, and γδT cell within cluster 2 represent “activated” cells. To explore this hypothesis in more detail, we examined the expression level of γδT cell activation and replication markers, namely, *Cd44, Cd69, Nr4a1*, and *Mki67*. We found that, upon infection, the γδT cells within cluster 2 robustly express *Cd69* and *Nr4a1*, and to a lesser extent *Mki67*, suggesting local activation and expansion of the γδT cells (**Figure 4D**). Spatial module scoring analysis identified that Vγ6^+^ cells localise mainly to the dermis and epidermis of naïve mice (**Figure 4E-F**), but their distribution changes in response to infection, and they localised to the subcutaneous adipose tissue (**Figure 4E-F**). Together, these results indicate that during infection, there is a population of Vγ6^+^ cells within the subcutaneous adipose tissue compartment that exhibits features of local activation. During cold challenge, Vγ6^+^ cells are known to drive thermogenesis and lipid mobilisation *via* lipolysis through crosstalk with brown and beige adipocytes^36, 37^. However, interactions between Vγ6^+^ cells and white adipocytes, in particular in the subcutaneous white adipose tissue during infection, has yet to be explored.

**Figure 4.**
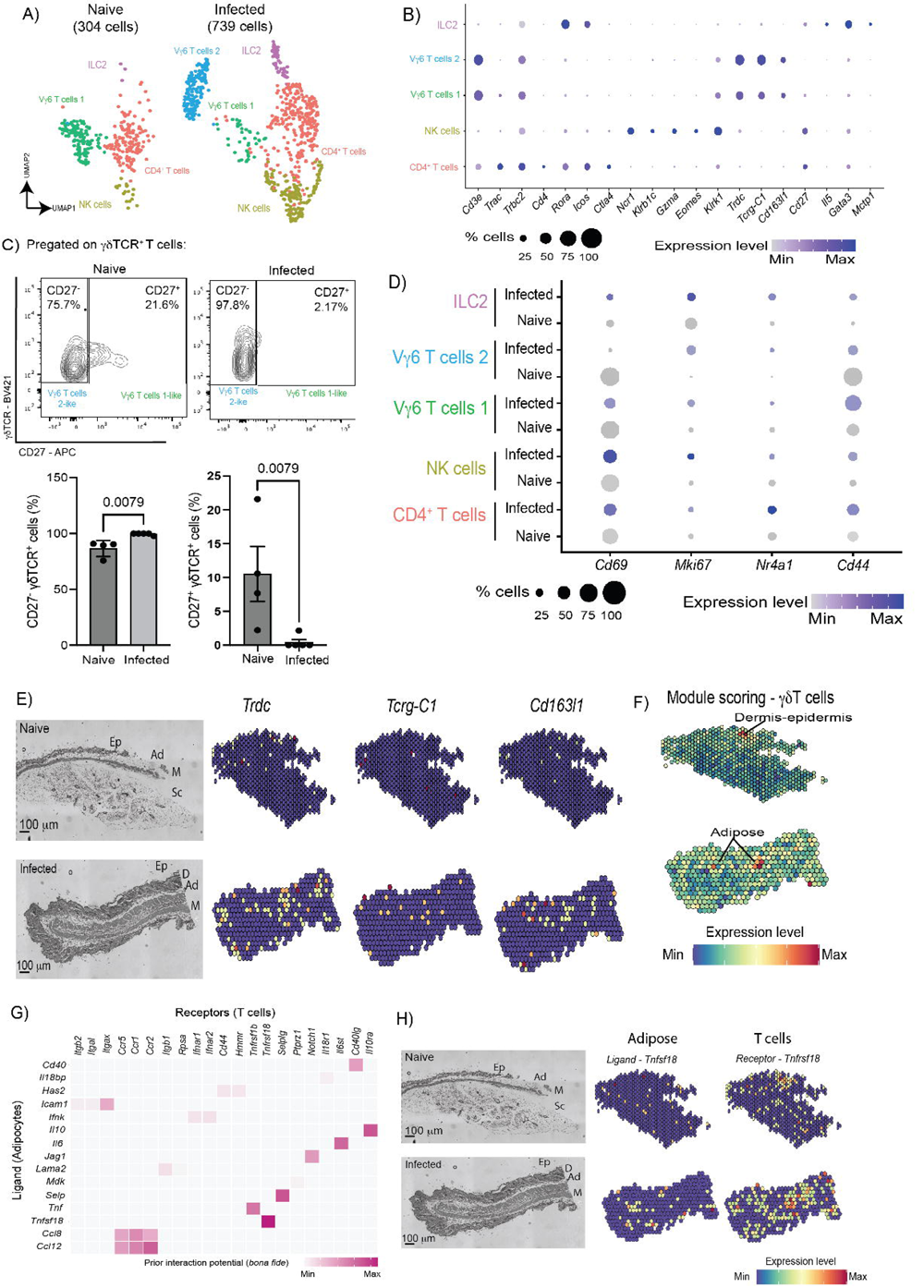
Chronic *T. brucei* infection triggers the activation of skin-dwelling Vγ6 γδT cells localised in the subcutaneous adipose tissue. **A)** Uniform manifold approximation and projection (UMAP) of 1,043 high-quality T cells from naïve (304 cells) and infected skin samples (739 cells). After reclustering, we detected a total of five subclusters including type 2 innate lymphoid cells (ILC2s), CD4^+^ T cells, NK cells, and γδT cells. **B)** Dot Plot depicting the expression level of top marker genes for the skin T cell subcluters. The dot size and the intensity of the colour represents the proportion of cells expressing the genes and the level of expression. **C)** Top panel: Representative flow cytometry analysis to determine the presence of CD27^-^ (IL-17-producing) and CD27^+^ (IFNg-producing) gdT cells in murine skin infected (*n* = 5 mice) with *T. brucei* and naïve controls (*n* = 4 mice). Bottom panel: Quantification of flow cytometry data. Statistical analysis was conducted using a parametric *T* test. A *p* value <0.05 was considered significant. **D)** Dot Plot depicting the expression level of top genes associated with T cell activation and TCR engagement for the skin T cell subcluters. The dot size and the intensity of the colour represents the proportion of cells expressing the genes and the level of expression. **E)** Spatial feature plot of canonical Vγ6^+^ cell marker genes. **F)** Spatial module scoring subclusters in the murine naïve and infected skin tissue for canonical γδ T cell marker genes. **G)** *In silico* cell-cell interaction analysis between adipocytes (“senders”) and T cells (“receivers”) based on the upregulation of ligand-receptor pairs. The heatmap is colour coded to represent the strength of the interaction. **H)** Spatial feature plot depicting the expression of adipose-derived ligand *Tnfsf18* and T-cell specific receptor*Tnfrsf18*, which is predicted to be one of the most robust interactions between these cell types during infection.

We next examined whether there is a cell-cell communication axis between adipocytes and all T cell clusters found in the skin during infection. NicheNet analysis indicates that adipocytes provide several critical cues for T cell recruitment and activation, including the activating factors *Cd40, Tnfsf18*, and *Icam1*, chemokines *Ccl12, Ccl8*, and cytokines *Tnf, Il10,* and *Il6* (**Figure 4G**). These adipocyte-derived ligands are predicted to be sensed by skin-resident T cells *via* the expression of *Cd40lg, Itgax, Il10ra, Il6st, Tnfrsf1b*, and *Tnfrsf18*. In the spatial context, we noted that the expression of adipocyte derived *Tnfsf18* is restricted to the subcutaneous adipose tissue in the infected skin, coinciding with the expression of T cells expressing their cognate receptor, *Tnfrsf18* (**Figure 4H**). Taken together, these results demonstrate that chronic skin infection with *T. brucei* leads to the local activation of a subpopulation of Vγ6^+^ cells in a process likely aided by subcutaneous adipocytes.

### Vγ6^+^ cells are essential for controlling skin inflammation, local CD8^+^ T cell activation, and subcutaneous adipose tissue wasting independently of skin-resident T_H_1 T cells

So far, our data indicates that Vγ6^+^ cells expand significantly in the chronically infected skin, and we predicted that these populations receive activation signals from adipocytes within the subcutaneous adipose tissue. To better understand if the skin-resident Vγ6^+^ cells are involved in a communication axis with subcutaneous adipocytes, we re-clustered the T cells and adipocytes together to conduct ligand-receptor mediated cell-cell communication analyses between these cell types. We found that Vγ6^+^ cells express several key genes involved in driving mesenchymal (including preadipocyte) differentiation and adipocyte lipolysis (**Figure 5A**). For instance, we detected an upregulation of the Cardiotrophin-like cytokine factor 1 (*Clcf1*) and Amphiregulin (*Areg*) in the Vγ6^+^ cells during infection, which are predicted to engage with their cognate receptors *Cntfr* and *Egfr*, respectively (**Figure 5A and B**). Other Vγ6^+^ cell-derived factors promoting mesenchymal differentiation are *Cd24a*, the Placental growth factor (*Pgf*), Sialophorin (*Spn*), and Pleiotrophin (*Ptn*), which are predicted to engage to *Pparg^+^* adipocytes *via* Selectin P (*Selp*), Neuropilin 1/2 *(Nrp1/Nrp2),* Syndecan 3 *(Sdc3)*, and Sialic Acid Binding Ig Like Lectin 1 (*Siglec1)*, respectively (**Figure 5A**). Interestingly, some of these Vγ6^+^ cell-derived factors are known to modulate thermogenesis, leading to mobilisation of lipids *via* lipolysis^36–38^. Together, this suggests that during infection Vγ6 γδT cells may be involved in promoting lipolysis to mobilise energy storage from adipocytes. Based on these observations, we hypothesised that, during infection, Vγ6^+^ cells expand in the skin and are important for driving subcutaneous adipose tissue wasting, limiting IFNγ-driven skin inflammation, and controlling parasite burden. To test our hypothesis, we infected mice with a double Vγ4/6 T cell knockout (on an FVB/N genetic background; Vγ4/6^-/-^) for a period of 25 days, and we monitored parasitaemia and clinical scores throughout infection. We observed that both Vγ4/6^-/-^ and FVB/N mice displayed the same levels of parasitaemia and parasite burden in the skin as measured by qRT-PCR against the trypanosome-specific *Prf2* gene (**S3 Figure**). We also detected both slender and stumpy forms in all dermal layers in both infected Vγ4/6^-/-^ and FVB/N mice without significant differences between strains (**S4 Figure**), suggesting that Vγ4^+^ and Vγ6^+^ cells are dispensable for controlling parasite burden in the skin.

**Figure 5.**
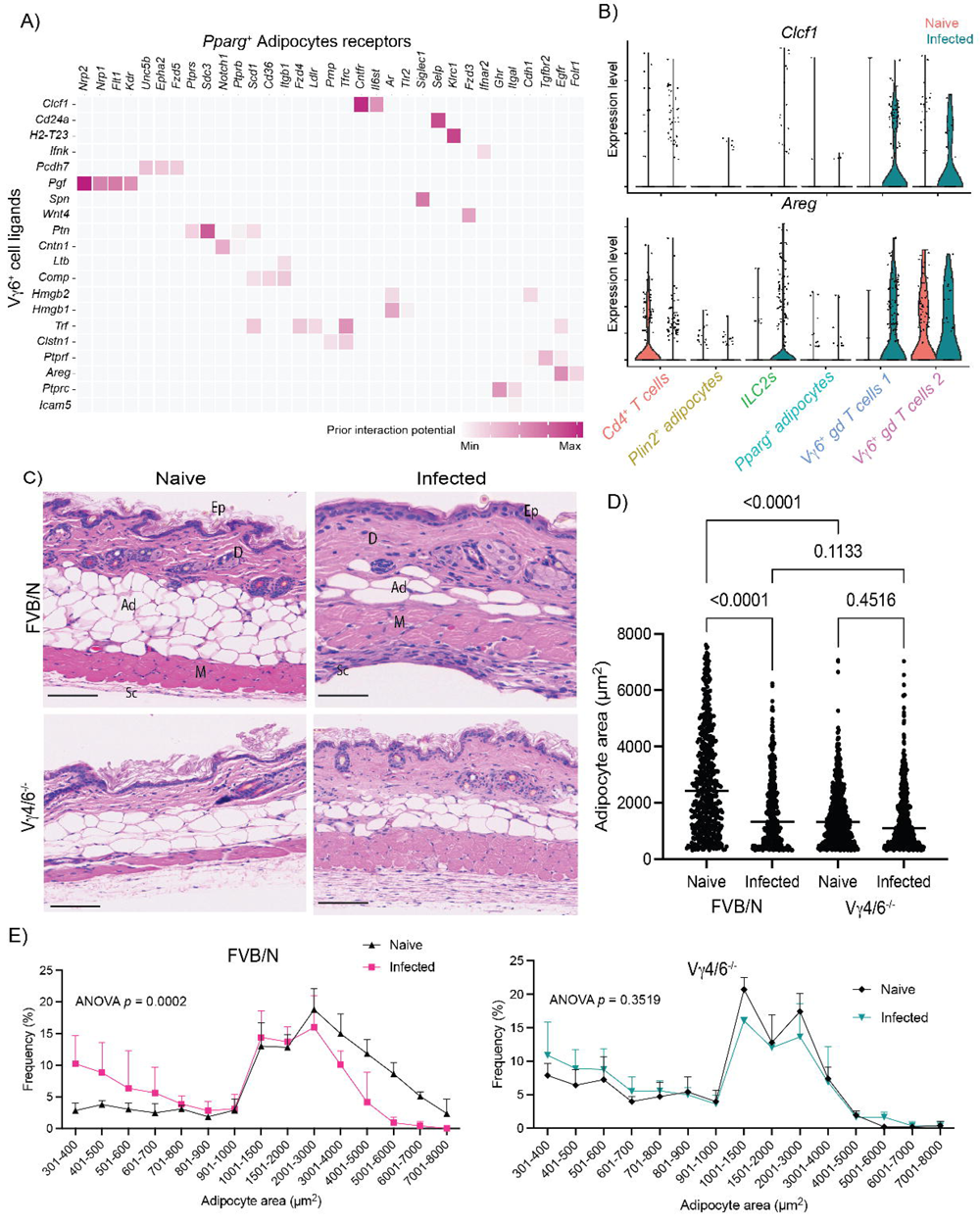
Vγ6^+^ cells are essential for controlling skin inflammation and subcutaneous adipose tissue wasting independently of skin-resident T_H_1 T cells. **A)** *In silico* cell-cell interaction analysis between Vγ6^+^ cells (“senders”) and *Pparg^+^* adipocytes (“receivers”) based on the upregulation of ligand-receptor pairs. The heatmap is colour coded to represent the strength of the interaction. **B)** Expression level of *Clcf1* and *Areg*, two of the most significant upregulated Vγ6^+^ cells-derived ligands predicted to interact with subcutaneous adipocytes. **C)** H&E staining from skin biopsies obtained from FVB/N and Vγ4/6^-/-^ naïve and infected mice. Scale bar: 100 μm. Ep, epidermis; D, dermis; Ad, adipose tissue; M, muscle; Sc, subcutis. **D)** Analysis of mean adipocyte area (µm^2^) in naïve and infected FVB/N and Vγ4/6^-/-^ mice. *N* = 5 biological replicates per group, from two independent experiments. Lipid droplets were measured from 3 distinct areas in each image and then combined for each biological replicate. A non-parametric, one-way ANOVA was used to determine the level of significant. A *p* value <0.05 is considered significant**. E)** Frequency plot of the adipocyte area represented in **(D)** for naïve and infected FVB/N (left panel) and Vγ4/6^-/-^ mice (right panel). *N* = 5 biological replicates per group, from two independent experiments. Lipid droplets were measured from 3 distinct areas in each image and then combined for each biological replicate. A non-parametric, one-way ANOVA was used to determine the level of significant. A *p* value <0.05 is considered significant.

We next examined histological sections of skin samples from infected Vγ4/6^-/-^ and FVB/N mice, as well as their counterpart naïve controls. Compared to infected FVB/N mice, the Vγ4/6^-/-^ mice displayed more severe signs of skin inflammation. Specifically, we observed higher follicular atrophy in the dermis and hypodermis, as well as diffuse lymphocyte aggregates containing large number of plasma cells and oedema in the subcutaneous adipose tissue compared to infected FVB/N mice (**Figure 5C and S3 Table**). Histological analysis indicated that, during infection, FVB/N mice lose a greater proportion of subcutaneous adipocytes than their naïve counterparts, consistent with previous reports in the trypanotolerant C57BL/6 background^24^ (**Figure 5C**). In contrast, infected Vγ4/6^-/-^ mice retain similar adipocyte numbers to their naïve counterparts (**Figure 5C and 5D**), highlighting a critical role for Vγ4/6 γδ T cells in modulating subcutaneous adipose tissue wasting. Morphometric analysis revealed that adipocytes in the Vγ4/6^-/-^ mice were significantly smaller in area in naïve animals compared to the FVB/N background (**Figure 5D**), potentially highlighting a role for the Vγ4^+^ and Vγ6^+^ cells in maintaining adipocyte function under homeostasis. Moreover, our morphometric analyses revealed that adipocytes within the subcutaneous adipose tissue of infected FVB/N mice were significantly smaller than those in naïve mice, whereas infection did not significantly impact adipocyte size in the Vγ4/6^-/-^ mice (**Figure 5E**).

Lastly, within the T cell compartment, we failed to detect significant differences in the frequency of skin-resident CD4^+^ and CD8^+^ T cells, or the frequency of IFNγ-producing CD4^+^ T cells (**S5 and S6 Figures**), indicating a dispensable role for Vγ4^+^ and/or Vγ6^+^ cells in limiting TH1-mediated T cell responses. However, we noted that in the skin of the Vγ4/6^-/-^ mice there was a significant increase in the frequency of IFNγ-producing CD8^+^ T cells compared to FVB/N controls, potentially suggesting that Vγ4^+^ and/or Vγ6^+^ cells may regulate the activation threshold of CD8^+^ T cells under homeostatic conditions (**S6 Figure**). Taken together, our results demonstrate that Vγ4^+^ and/or Vγ6^+^ cells are critical for limiting skin pathology, at least in part by controlling the activation state of skin-resident IFNγ^+^ CD8^+^ T cell, as well driving subcutaneous adipose tissue wasting, in a process likely involving IL-17 signalling.

## Discussion

To address key questions about the immune response of the skin to chronic infection with *T. brucei*, this study aimed to characterise changes in skin cell populations using single cell transcriptomics, as well as to determine how the cell populations detected by single cell transcriptomics are distributed throughout the skin during infection using spatial transcriptomics. With this information, we then modelled cell-cell interactions in the skin during *T. brucei* infection, to understand immune-stromal crosstalk and how this influences the immune response to infection. Here, using a combination of cutting-edge technologies and genetic murine models, we demonstrated that IL-17-producing Vγ6^+^ cells play a critical role in the controlling skin inflammation (**Figure 6**). Furthermore, our data highlight a previously unappreciated interaction between subcutaneous interstitial preadipocytes and mature adipocytes and skin-dwelling T cells (including γδ T cells), which mediates T cell responses and subcutaneous adipose tissue wasting (**Figure 6**). We first generated a spatially resolved single cell atlas of the murine skin during chronic *T. brucei* infection. From these analyses, several observations are worth discussing in detail. First, we observed significant changes in the skin stromal and immune compartment without the formation of granulomatous lesions, indicative of subclinical inflammatory processes when compared to naïve controls. Using module scoring of inflammatory cytokines and chemokines, as well as genes associated with antigen presentation, we detected inflammatory signatures predominantly in populations of *Cd14*^+^ monocytes, Langerhans cells, and interstitial preadipocytes located in the subcutaneous adipose tissue.

**Figure 6.**
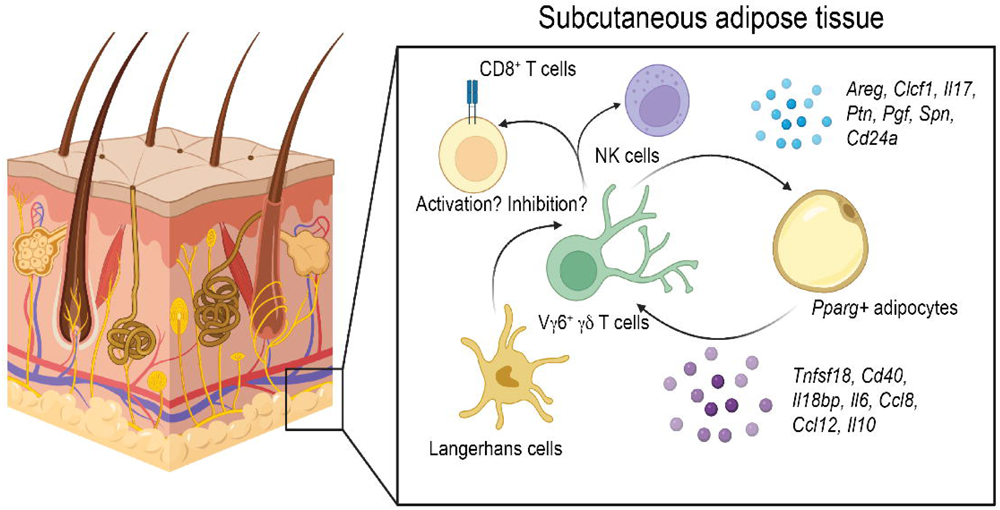
Proposed model of stromal-immune interactions in the skin during *T. brucei* infection. Based on our spatially-resolved single cell atlas, we propose a model whereby Vγ6^+^ cells act concertedly with *Pparg^+^* adipocytes (and potentially preadipocytes) to coordinate local immune responses, either *via* the recruitment of immune cells (e.g., CD8^+^ T cells and NK cells). The *Pparg^+^* adipocytes in this context provide important cues for T cell activation that we hypothesise might be involved in triggering Vγ6^+^ cells-mediated responses. Other stromal cells such as Langerhans cells and keratinocytes are also likely to be involved in this process via antigenic presentation.

Following a reclustering of stromal cells, we identified two populations of *Dpp4*^+^ interstitial preadipocytes (IPA1 and IPA2) that upregulate inflammatory cytokines, chemokines, and molecules associated with antigen presentation. The chemokines *Cxcl1*, *Cxcl9* and *Cxcl10* were upregulated in both populations of interstitial preadipocytes but were higher in IPA2. These chemokines are secreted to recruit neutrophils^39^, CD4^+^ and γδ T cells^40, 41^, and natural killer cells^42^, and their upregulation exclusively by preadipocytes suggests that these cells are critical drivers of immune recruitment to the skin during *T. brucei* infection. Supporting this, in the skin of infected mice we found expansion of CD4^+^ T cells, γδ T cells, and NK cells, and interestingly this may represent a feedback loop whereby these immune cells suppress differentiation of preadipocytes to mature adipocytes, as observed using *in vitro* models^43^. In addition to preadipocytes, we also found that mature adipocytes upregulate chemokines during infection, including *Ccl8* and *Ccl12*, which are drivers of monocyte recruitment^44^. Moreover, these cells upregulated *Il6* and *Il10*, which were predicted to communicate with T cells through *Il6st* and *Il10ra*, respectively. To our knowledge, although adipose tissue immune populations are known to express IL-10^45^, and adipocytes express the IL-10 receptor^46^, this is the first time that adipocyte *Il10* expression has been reported. In the subcutaneous adipose tissue, IL-10 signalling limits energy expenditure and lipolysis in mouse models of cold exposure and obesity^46^, but its effects on adipocytes during infection remain unknown. Our observations may suggest that during *T. brucei* infection, IL-10 acts in both an autocrine and paracrine fashion, whereby adipocytes secrete the cytokine and it suppresses adipose tissue lipolysis, whilst simultaneously suppressing CD4^+^ T cell activity^47^. Conversely, IL-6 is a driver of lipolysis and fatty acid oxidation^48^ and is associated with weight loss and fat wasting in diseases such as HIV^49^ and cancer^50^. It is, therefore, unclear how these two cytokines impact adipocyte activity when both are present.

Importantly, we also identified a population of skin-resident Vγ6^+^ T cells that expand in response to skin infection. These Vγ6^+^ cells are primed to produce IL-17A and IL-17F during development in the thymus^51, 52^, and upon maturation they migrate to multiple tissues throughout the body, including the skin, where they become resident immune cells, offering a first line of response to infection^53^. The γδ T cells that we identified in our dataset express markers putatively associated with activation, including *Cd44, Cd69*, and *Nr4a1*, potentially suggesting the existence of local drivers of γδ T cell activation in the infected skin. Both *in silico* predictions and *in vivo* analyses indicate that these γδ T cells are likely to be IL-17A-producing Vγ6^+^ cells based on the expression levels of *Tcrg-C1*, *Cd163l1*, and CD27, as previously reported^35^. We also observed two populations of Vγ6^+^ cells, which we hypothesise represent “resting” and “activated” populations, based on expression of *Cd44, Cd69*, and *Nr4a1*. However, unlike other γδ T cell populations in the skin, such as dendritic epidermal T cells that reside solely in the dermis, Vγ6^+^ cells may be able to recirculate between tissues^54, 55^. In this scenario, our data may provide evidence of one Vγ6^+^ population exiting the skin and a separate population entering the skin during infection. However, future studies are required to dissect the migratory dynamics of skin Vγ6^+^ cells in the context of skin infection.

Our findings further highlight the importance of IL-17A signalling in the skin of mice infected with *T. brucei*, consistent with our previous work proposing IL-17A as a critical driver of subcutaneous and inguinal adipose tissue wasting^24^. Interestingly, in the spatial context, these IL-17A-producing Vγ6^+^ cells are located in the subcutaneous adipose tissue layer of the skin, and are predicted to establish crosstalk with adipocytes via several molecules, including T cell co-stimulatory signals such as *Tnfsf18*, which engages with GITR (*Tnfrsf18*) to lower the T cell activation threshold^56^. These cells also express *Cd40*, indicating a previously unappreciated crosstalk between stromal adipocytes and Vγ6^+^ cells during *T. brucei* infection in the skin. Consistent with this, mice lacking Vγ4/6^+^ T cells display a higher number of plasma cells and more severe skin inflammation compared to wild type controls. Mice deficient in Vγ4^+^ and Vγ6^+^ are known to develop increased numbers of plasma cells and spontaneous germinal centre formation^57^, which may dysregulate the immune response to infection. We found that the increased inflammation in the skin of infected Vγ4/6^-/-^ mice may be in part mediated by controlling the activation threshold of skin-resident CD8^+^ T cells. The increased capacity of CD8^+^ T cells to produce IFNγ in the skin of naïve Vγ4/6^-/-^ mice suggests that these cells play a role in constraining CD8^+^ T cell activity under homeostasis. An alternative possibility, is that the increased severity of skin inflammation in infected Vγ4/6^-/-^ mice is due to exacerbated recruitment of neutrophils, as reported in metastatic breast cancer^58^. Strikingly, we found that Vγ4/6^-/-^ mice do not lose subcutaneous adipose tissue to the same extent as wild type controls during infection, mirroring our previous studies where we proposed IL-17A as a driver of infection-associated adipose tissue wasting^24^. In both wild type and Vγ4/6^-/-^ mice we observed comparable parasite tissue burden, with both slender and stumpy developmental forms of the parasite readily detected in all the dermal layers of these mice. Furthermore, we failed to detect significant differences in the frequency of skin-resident T_H_1 T cells, which strongly suggest that both parasites and T_H_1 T cells are dispensable for driving subcutaneous adipose tissue wasting, which is typically observed in this experimental infection setting^24, 33^. Thus, it is tempting to speculate that IL-17A-producing Vγ6^+^ cells (and potentially other sources of IL-17A such as TH17 T cells) promote subcutaneous adipose tissue lipolysis to fuel an efficient immune response against the parasites, although the molecular mechanisms underlying this process need to be investigated in more detail. However, in the study presented here we were unable to dissect the relative contribution of Vγ4^+^ or Vγ6^+^ cells to skin inflammation, or the interactions between different γδ T cell subsets that reside in the skin under both homeostatic conditions and inflammation.

Our data strongly indicate that the subcutaneous adipose tissue is an active site for immune priming and activation, placing the adipocytes at the core of this process. Thus, we propose a model whereby subcutaneous adipocytes (in addition to Langerhans cells and keratinocytes) have a critical role as coordinators of local innate and adaptive immune responses. In the context of trypanosome infection, subcutaneous adipocytes may detect the presence of parasites (e.g., *via* Toll-like receptor signalling) to trigger the recruitment and activation of innate immune cells such as γδ T cells to mobilise energy stores to meet the energetic requirements needed to control infection, as recently proposed^59^.

Together, our spatially resolved atlas of the murine skin during *T. brucei* infection offers a resource to the community interested in understanding how chronic infections affect skin homeostasis and immunity. Future work is required to examine the consequences of infection on adipocyte differentiation and function, and whether these processes are directly controlled by γδ T cells, as shown during cold exposure^36, 37^. Furthermore, our dataset provides strong evidence for an engagement of several cell types within the stromal compartment in the skin in response to infection, and may suggest that the efficacy, robustness, and timing of the local immune response may be determined by cell types traditionally associated with non-immunological functions such as mesenchymal cells (including interstitial preadipocytes). We envision that future work dissecting the role of these various cell types and communication axis will address some of the fundamental questions arising from this study.

## Acknowledgments and funding

We thank Julie Galbraith and Pawel Herzyk (Glasgow Polyomics, University of Glasgow) for their support with library preparation and sequencing. Similarly, we would like to thank the technical staff at the University of Glasgow Biological Services for their assistance in maintaining optimal husbandry conditions and comfort for the animals used in this study. We thank the Maria Kasper Lab, Karolinska Institutet, Sweden for their advice on single cell skin dissociation. We thank Dr Jean Rodgers for the work conducted under her Home Office Animal License (PPL No. PC8C3B25C). This work was funded by a Sir Henry Wellcome postdoctoral fellowship (221640/Z/20/Z to JFQ), a Wellcome Trust FutureScope grant (104111/Z/14/Z Wellcome Centre for Integrative Parasitology to JFQ), and a Wellcome Trust Senior Research fellow (209511/Z/17/Z to AML). SBC is supported by a grant from the Annie McNab Bequest (CRUK Beatson Institute), Breast Cancer Now (2018JulPR1101, 2019DecPhD1349, 2019DecPR1424), Cancer Research Institute (CLIP award), Cancer Research UK (RCCCEA-Nov21\100003, EDDPGM-Nov21\100001, DRCNPG-Jun22\100007), and Pancreatic Cancer Research and Worldwide Cancer Research (22-0135). PC and MCS are supported by a Wellcome Trust Senior Research fellowship (209511/Z/17/Z) awarded to AML. RH is a Wellcome Trust PhD student (Wellcome Trust 218518/Z/19/Z). The authors declare that the research was conducted in the absence of any commercial or financial relationships that could be construed as a potential conflict of interest.

## Author contributions

**Conceptualisation:** JFQ, MCS. **Methodology:** JFQ, MCS, PC, RH, JO, BC, AL, AC, NK, SBC, AML. **Formal analysis**: JFQ, MCS, PC, RH, JO, BC, AL. **Writing – original draft:** JFQ, MCS**. Writing – reviewing and editing:** JFQ, MCS, PC, RH, JO, BC, AL, AC, SBC, NK, AML. **Funding acquisition:** JFQ, AML. The authors declare that they have no competing interests, commercial or otherwise. Correspondence and requests for materials should be addressed to Annette MacLeod or Juan F. Quintana

## Supplementary Figures

**Supplementary figure 1.**
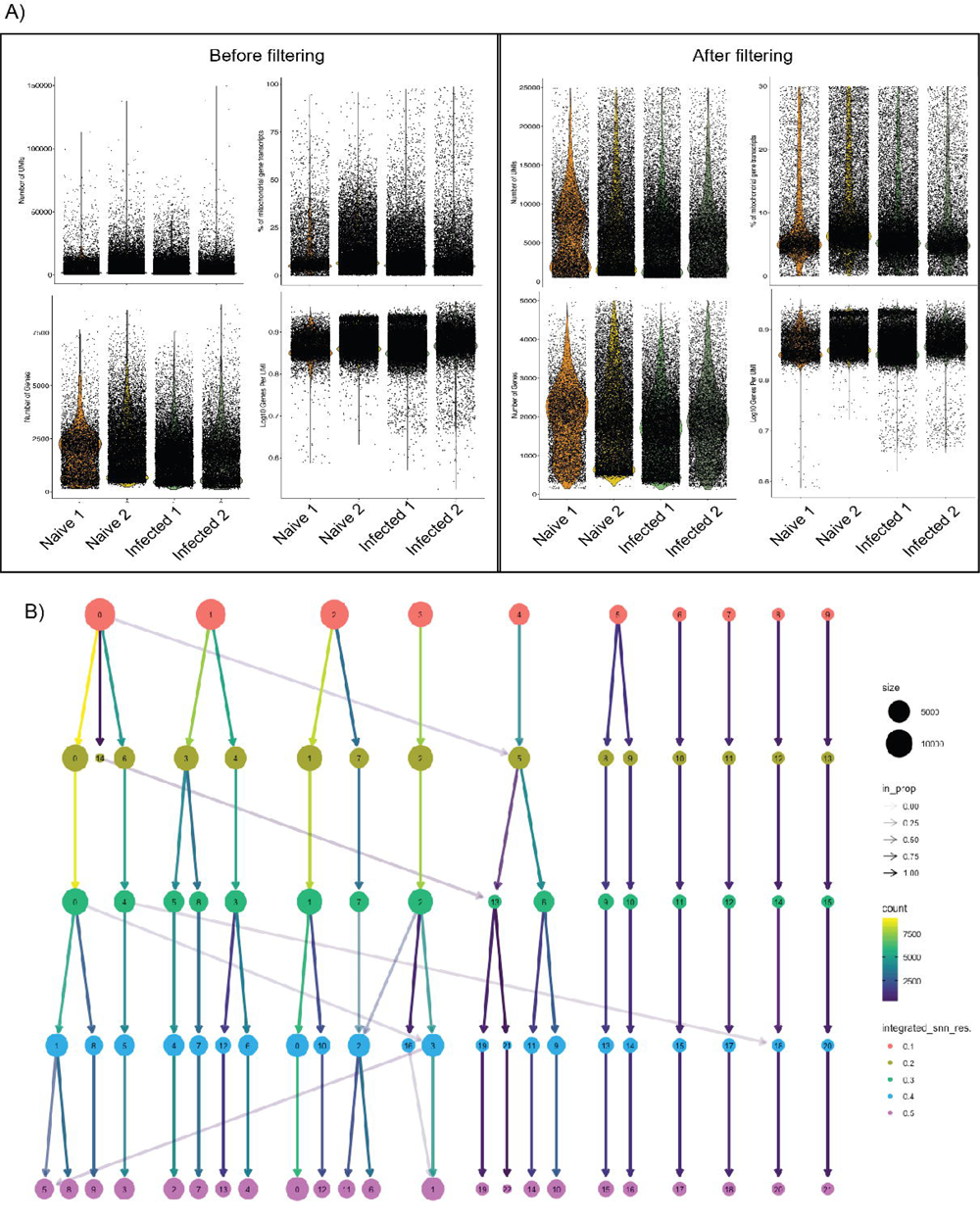
Quality control measurements of the murine single cell transcriptomics dataset. **A)** Number of Unique molecular identifies (UMIs), genes, mitochondrial reads, and library complexity (Log10 UMIs/gene) before (left panel) and after (right panel) applying filtering parameters. **B)** Clustree output representing the relationship between different cell clusters at various levels of resolution using the function *FindClusters*.

**Supplementary figure 2.**
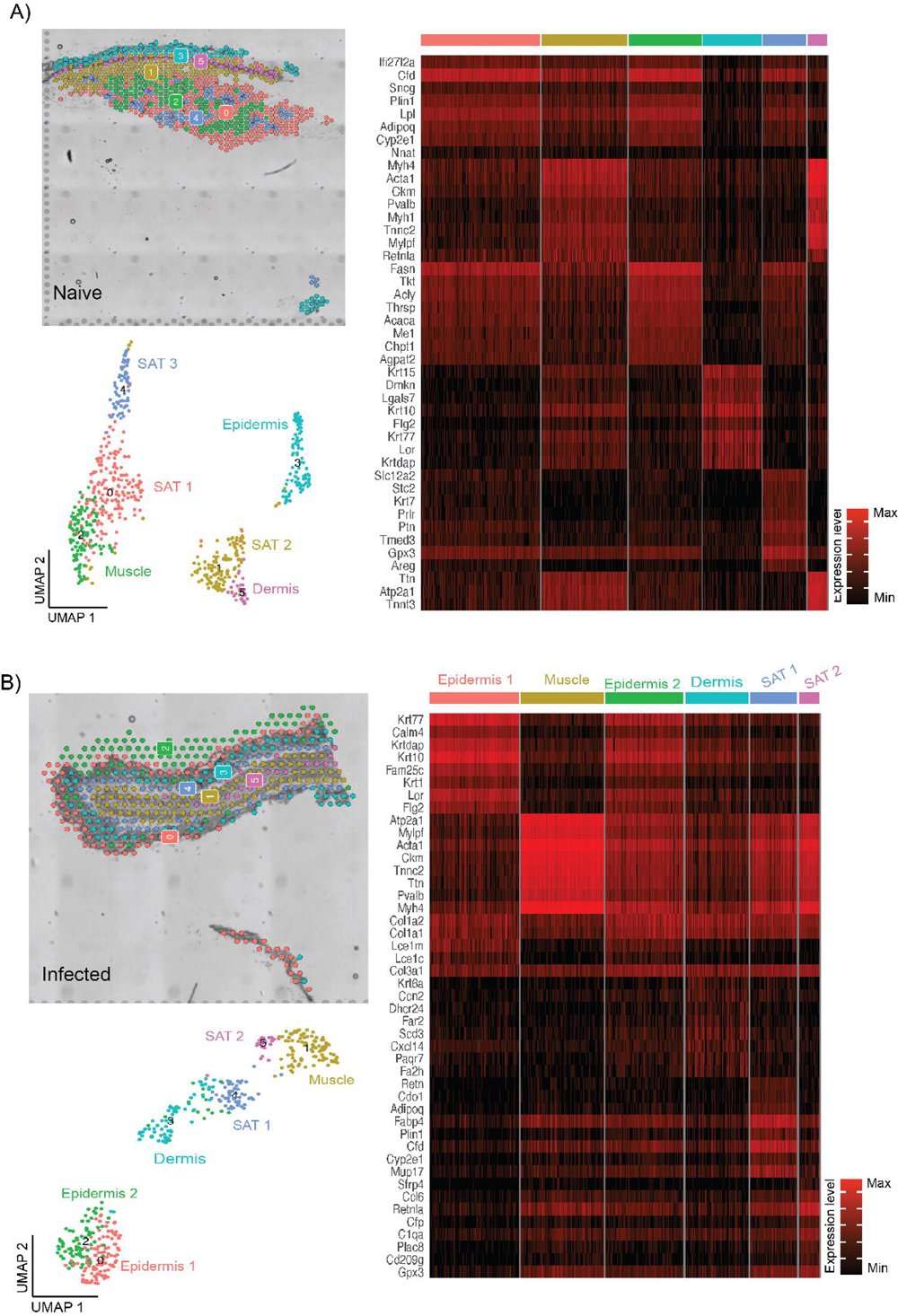
Quality control of 10X Visium datasets from the mouse skin over the course of infection with *T. brucei*. Spatial clusters and marker genes for each spatial cluster in the naïve **(A)** and infected **(B)** murine skin using 10X Visium spatial transcriptomics.

**Supplementary figure 3.**
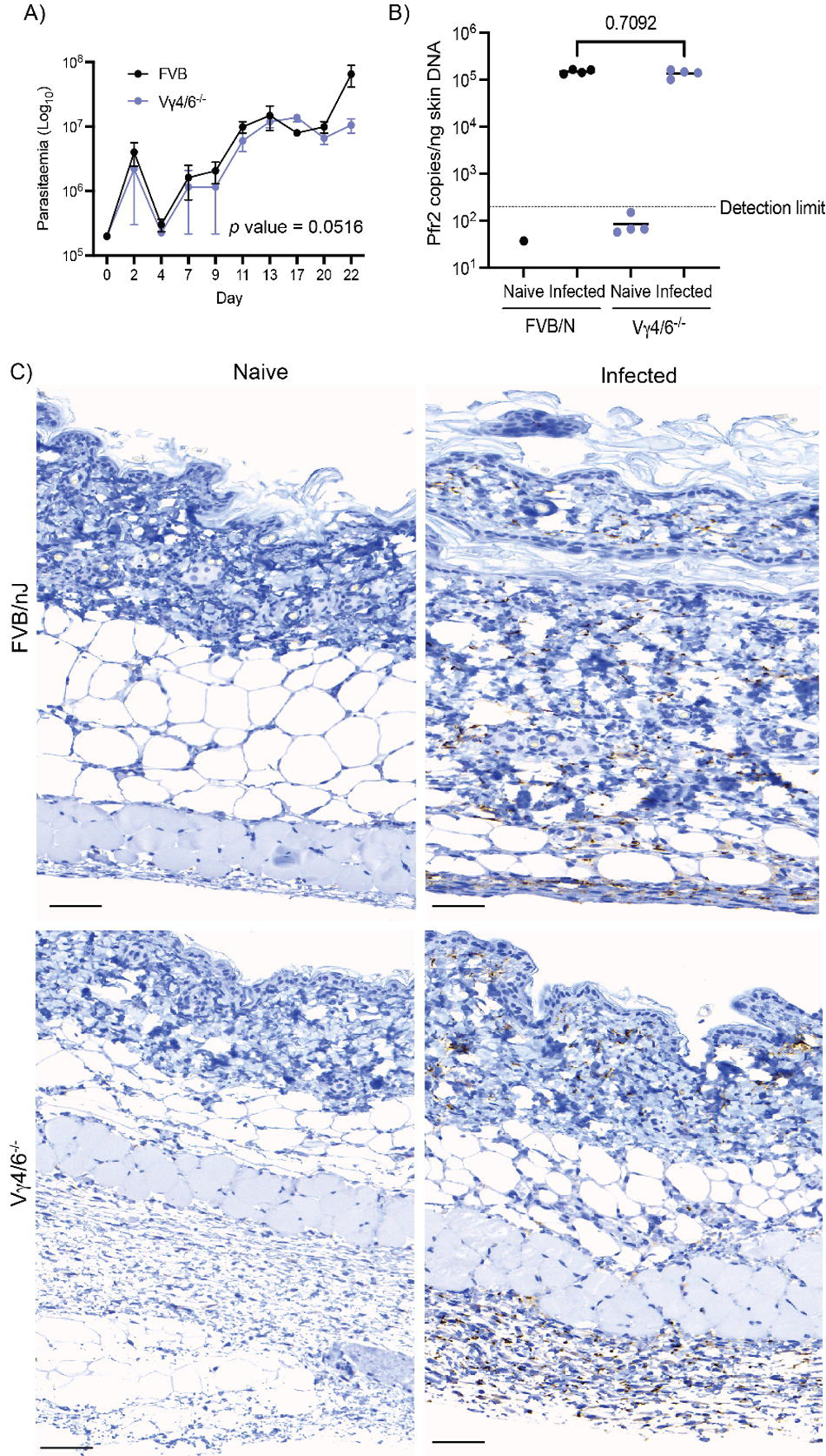
Characterisation of the Vγ4/6^-/-^ mice during *T. brucei* infection. **A)** Measurement of circulating parasitaemia in both female FVB/N (*n* = 4) and Vγ4/6^-/-^ mice (*n* = 4) for a period of 22 days. ANOVA test with multiple corrections. A *p* value <0.05 is considered significant. **B)** qRT-PCR of skin-dwelling trypanosomes by measuring the trypanosome-specific gene *Prf2* by qRT-PCR. The *Prf2* copy numbers were normalised per ng of skin DNA from naïve and infected FVB/N and Vγ4/6^-/-^ mice (*n* = 4 mice/group). *T* test comparison between infected samples. A p value <0.05 is considered significant. The detection limit, as measured in skin biopsies from naïve controls, is also indicated with a dotted line. **C)** Representative immunohistochemistry of murine skin biopsies from naïve and infected FVB/N and Vγ4/6^-/-^ mice to detect the presence of *T. brucei* using an antibody against the trypanosome-specific luminal binding protein 1 (BiP). Scale bar = 50 μm.

**Supplementary figure 4.**
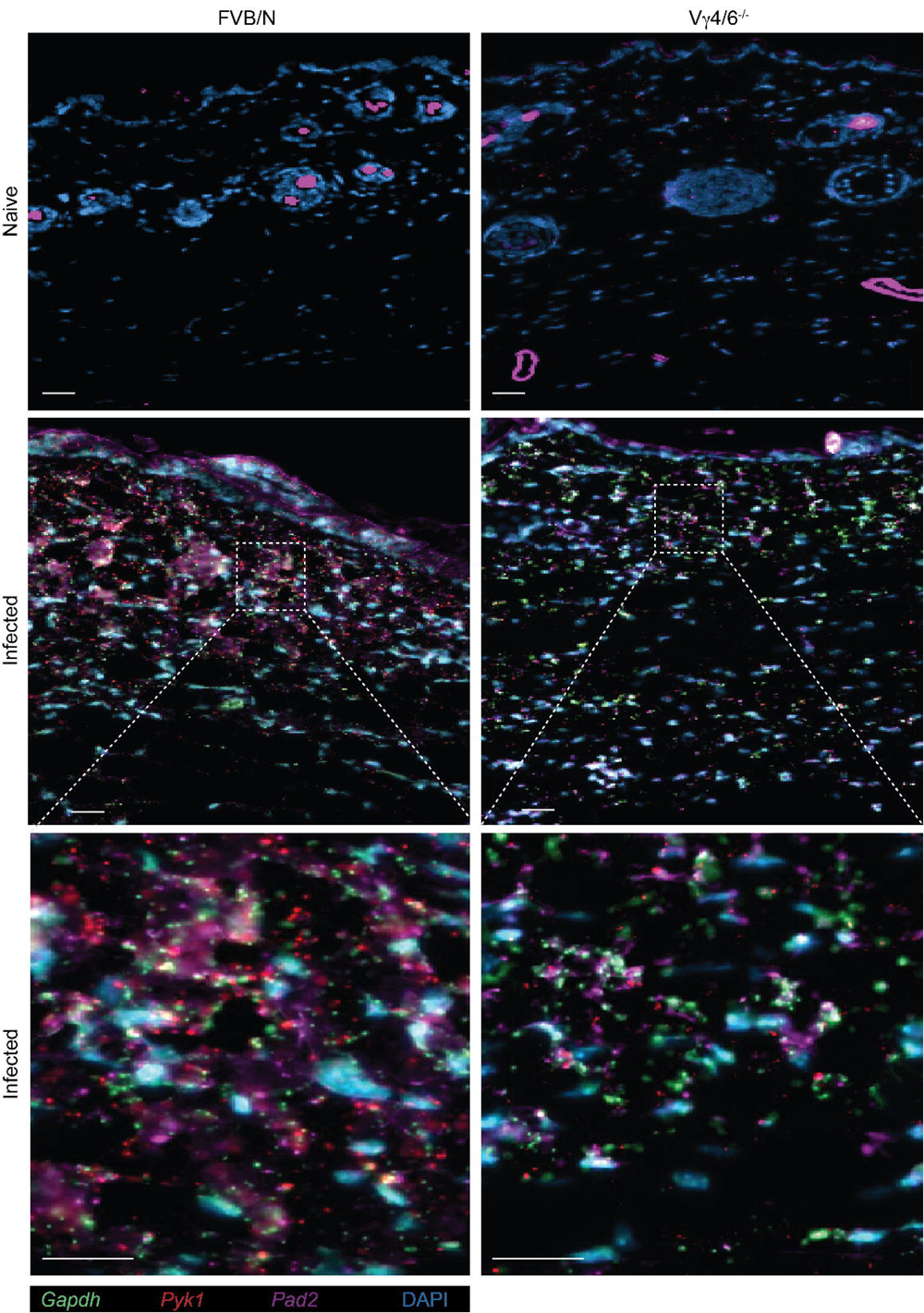
Single molecule fluorescent *in situ* hybridisation (smFISH) analysis showing the spatial distribution of *T. brucei* developmental stages in the skin of FVB/N and Vγ4/6^-/-^ mice. **Top panels:** skin sections from naïve FVB/N and Vγ4/6^-/-^ mice. **Middle panels:** skin sections from infected FVB/N and Vγ4/6^-/-^ mice depicting the expression of the *T. brucei*-specific transcripts *Gapdh* and *Pyk1* (slender specific markers) and *Pad2* (stumpy specific marker). The dotted square indicates an area selected for magnification. **Bottom panels:** Magnified fields of the infected samples showing the distribution of both slender and stumpy markers in the skin of FVB/N and Vγ4/6^-/-^ mice. Scale bar: 50 μm.

**Supplementary figure 5.**
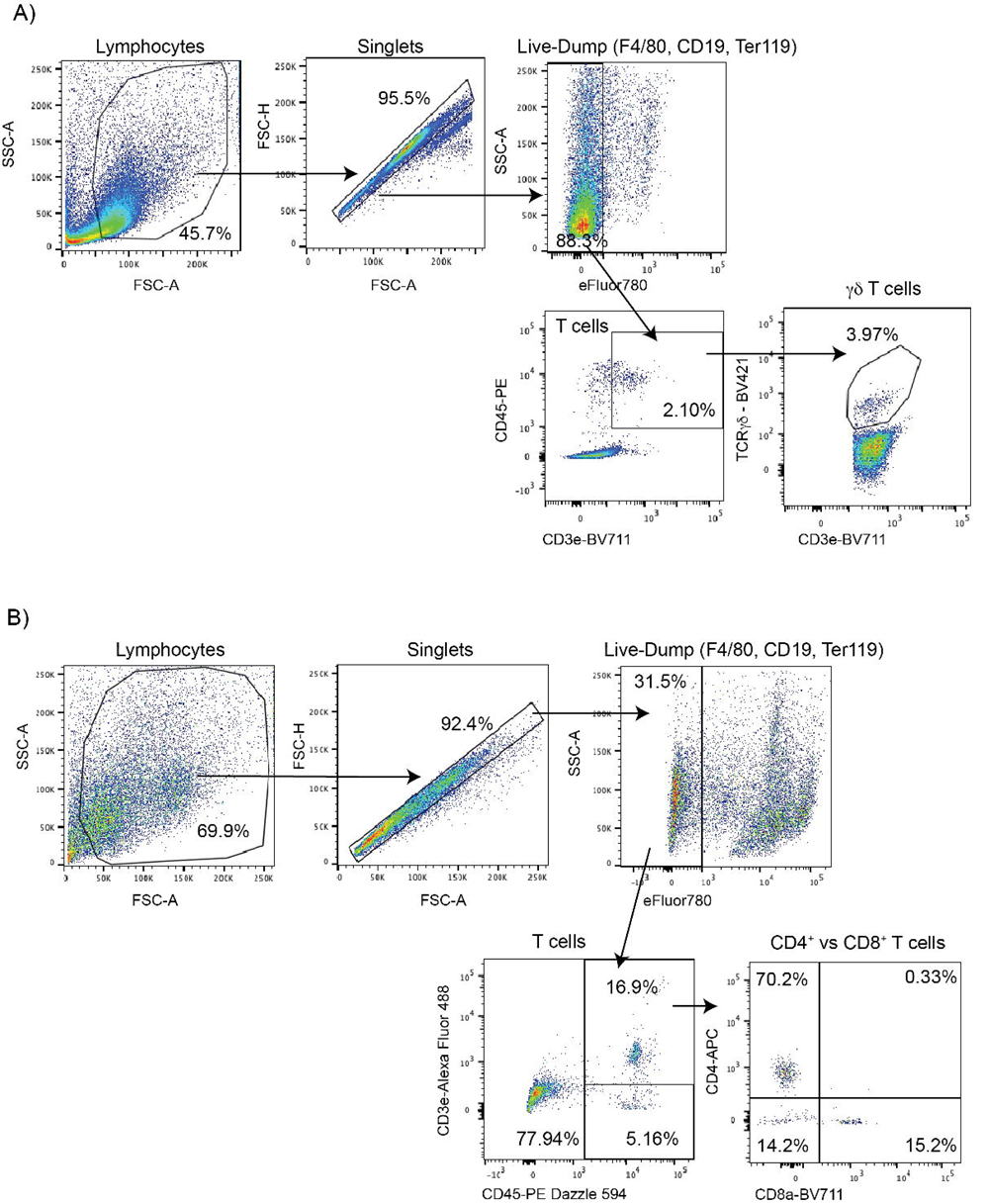
Flow cytometry analysis of skin-resident T lymphocytes. Gating strategy for the identification of skin γδ T cells **(A)** and CD4^+^ and CD8^+^ T cells **(B)**.

**Supplementary figure 6.**
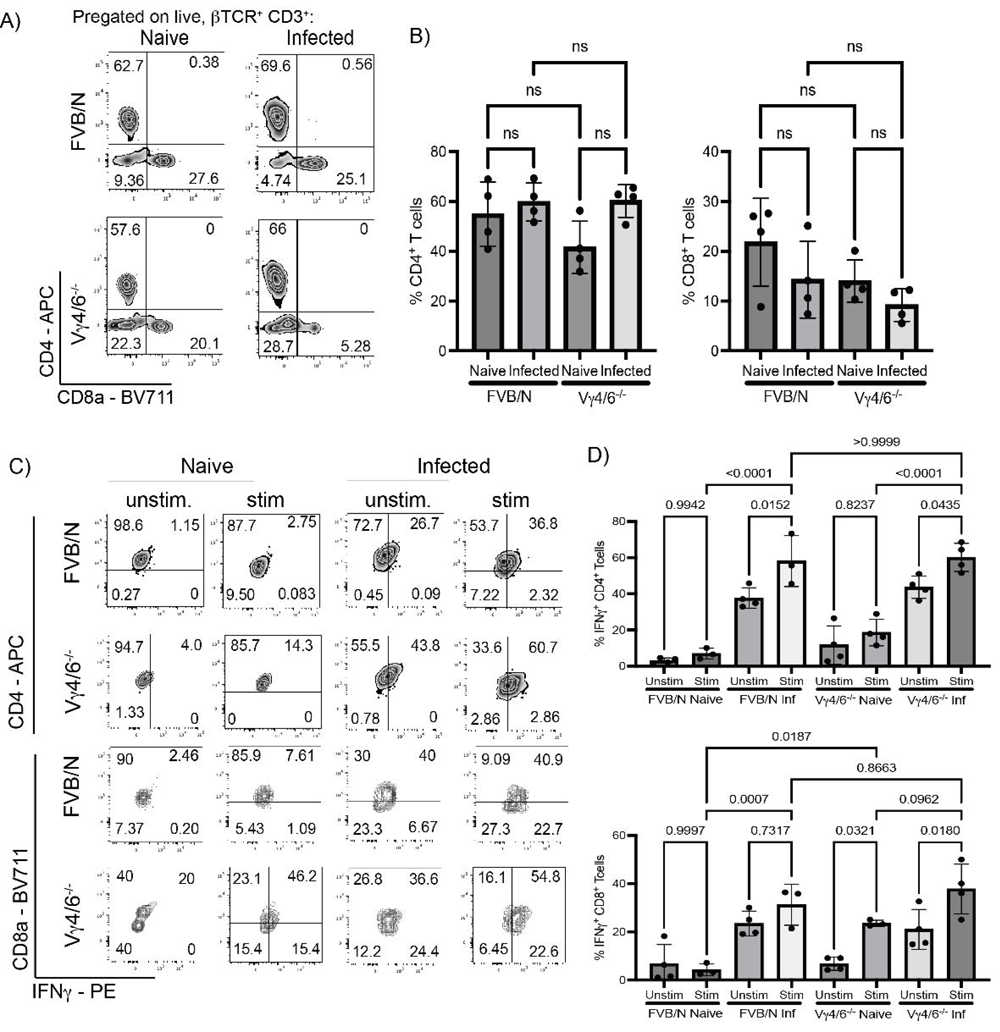
Quantification of skin-resident lymphocytes in the Vγ4/6^-/-^ mice during *T. brucei* infection. **A)** Representative flow cytometry analysis of skin CD4^+^ and CD8^+^ T cells in naïve and infected FVB/N and Vγ4/6^-/-^ mice (*n* = 4 mice/group). **B)** Quantification of the frequency of skin CD4^+^ (left panel) and CD8^+^ T cells (right panel) in naïve and infected FVB/N and Vγ4/6^-/-^ mice (*n* = 4 mice/group) as shown in (C). ANOVA test with multiple corrections. A *p* value <0.05 is considered significant. **C)** Representative flow cytometry analysis of *ex vivo* recall assay to determine the production of IFNγ in skin-resident CD4^+^ and CD8^+^ T cells in naïve and infected FVB/N and Vγ4/6^-/-^ mice (*n* = 4 mice/group). **D)** Quantification of the frequency of skin CD4^+^ (top panel) and CD8^+^ T cells (bottom panel) in naïve and infected FVB/N and Vγ4/6^-/-^ mice (*n* = 4 mice/group) as shown in (C).

## Table legend

**Supplementary table 1. Overview of the mouse skin single cell transcriptomics during chronic *T. brucei* infection.** **S1A)** Quality control including mean reads per cell and median genes per cell before and after filtering out low quality cell types. **S1B)** Overview of the major cell types detected in the single cell dataset at a resolution of 0.4. The marker genes are also included. **S1C)** Overview of the stromal cells detected in the skin dataset at a resolution of 0.3. The marker genes for these clusters, as well as representative UMAP plots are also included. **S1D)** As in S1C, but for the myeloid cells at a resolution of 0.3**. S1E)** As in S1C, but for the T cells at a resolution of 0.3.

**Supplementary table 2. Overview of the spatial transcriptomics of the mouse skin during chronic *T. brucei* infection.** **S2A)** Overview of the spatial transcriptomics project, including total number of reads sequenced per biological replicate, the median number of genes per spot and the percentage of mappable reads to the mouse genome (mm10). **S2B)** Mouse marker genes identified in the 10X Visium spatial transcriptomics datasets.

**Supplementary table 3. Histopathological analysis of biopsies taken from naïve and infected FVB/NJ and Vγ4/6^-/-^ mice.** The skin biopsies were harvested at 21 days post-infection (*n* = 4 mice/group), fixed in 10% PFA and counterstained with the *T. brucei*-specific antibody TbBiP. Uninfected animals (*n* = 4) were included as naïve controls. Histological examinations were scored using H&E staining and BiP staining and was conducted double-blinded. The results reported in this table are in comparison to naïve controls for the corresponding genetic background, and encompass a detailed analysis of the epidermis, dermis, hypodermis, skeletal muscle, and the subcutaneous adipose tissue. The column labelled “BiP” (columns AD to AI) represents the results from the immunohistochemistry analysis against the anti-Trypanosoma anti-BiP antibody.

